# Reconstitution of the *S. aureus agr* quorum sensing pathway reveals a direct role for the integral membrane protease MroQ in pheromone biosynthesis

**DOI:** 10.1101/2021.12.29.473670

**Authors:** Aishan Zhao, Steven P. Bodine, Qian Xie, Boyuan Wang, Geeta Ram, Richard P. Novick, Tom W. Muir

## Abstract

In *Staphylococcus aureus*, virulence is under the control of a quorum sensing (QS) circuit encoded in the accessory gene regulator (*agr*) genomic locus. Key to this pathogenic behavior is the production and signaling activity of a secreted pheromone, the autoinducing peptide (AIP), generated following the ribosomal synthesis and post-translational modification of a precursor polypeptide, AgrD, through two discrete cleavage steps. The integral membrane protease AgrB is known to catalyze the first processing event, generating the AIP biosynthetic intermediate, AgrD (1-32) thiolactone. However, the identity of the second protease in this biosynthetic pathway, which removes an N-terminal leader sequence, has remained ambiguous. Here, we show that MroQ, an integral membrane protease recently implicated in the *agr* response, is directly involved in AIP production. Genetic complementation and biochemical experiments reveal that MroQ proteolytic activity is required for AIP biosynthesis in *agr* specifiy groups -I and -II, but not group-III. Notably, as part of this effort, the biosynthesis and AIP-sensing arms of the QS circuit were reconstituted together *in vitro*. Our experiments also reveal the molecular features guiding MroQ cleavage activity, a critical factor in defining *agr* specificity group identity. Collectively, our study adds to the molecular understanding of the *agr* response and *Staphylococcus aureus* virulence.

## Introduction

Quorum sensing (QS) is a common form of chemical communication used by bacteria to coordinate behavior.^1^ Small molecules—acting as chemical signals—are continuously produced, secreted, and recognized by bacteria to dynamically monitor local population density and direct gene expression. In *Staphylococcus aureus*, a common pathogenic Gram-positive bacterium widely associated with community acquired and nosocomial infections, a QS system helps optimize bacterial behaviors throughout all stages of infection, from colonization of infected hosts to virulence factor expression and resultant pathogenicity.^2, 3, 4, 5, 6, 7^ Pharmacological control of the *S. aureus* QS system would allow for the modulation of bacterial behavior, including attenuation of the detrimental effects of *S. aureus* infection, without imposing selective pressure to trigger the development of antibiotic resistance.^8, 9^ This makes the *S. aureus* QS circuit an attractive drug target and has motivated intense study of this system over the last three decades.^2,7, 10^

Much of the biochemical machinery necessary for *S. aureus* QS is encoded by the accessory gene regulator (*agr*) operon, which contains four open reading frames, *agrBDCA*.^2^ Biochemical studies have assigned specific function to each of these gene products, leading to a detailed mechanistic understanding of much of the system.^11, 12, 13, 14, 15, 16, 17^ AgrC is a transmembrane histidine phosphokinase signal receptor which is acted upon by a peptide pheromone, the *agr* auto-inducing peptide, AIP, of which AgrD is the precursor.^18, 19^ The AIP, a 5-membered macrocycle with a 2-4 amino acid N-terminal tail, is processed in two steps, of which the first is an AgrB-catalyzed cleavage of the C-terminal 18 amino acids concomitantly with the AgrB-induced formation of a thiolactone bond between the C-terminal carboxyl and an internal cysteine, generating a biosynthetic intermediate, the AgrD (1-32) thiolactone (**Figure 1A**).^15^ The N-terminal amphipathic helical domain of the thiolactone intermediate is cleaved in a second, separate proteolytic event, releasing the mature AIP into the extracellular milieu to serve as an index of local bacterial population density. When the AIP concentration exceeds a threshold, it is recognized by the sensor domain of the receptor histidine kinase, AgrC, leading to structural changes in the AgrC cytosolic domain that activates autophosphorylation and subsequent phosphoryl-transfer to the response regulator, AgrA.^14, 16, 17^ Together, AgrC and AgrA form a two-component signaling (TCS) pathway that up-regulates the transcription of the *agr* operon, leading to a positive feedback loop of AIP production and recognition.^7, 18^ Activated AgrA also promotes the transcription of the multifunctional RNA, RNAIII, which serves as the message for expression of the virulence factor δ-hemolysin and as a regulatory RNA suppressing the expression of adhesins and promoting the expression of various other virulence factors.^20, 21^ By this mechanism, *agr* QS functions as a master regulator of *S. aureus* biofilm dispersal and virulence.^7, 22^

**Figure 1.**
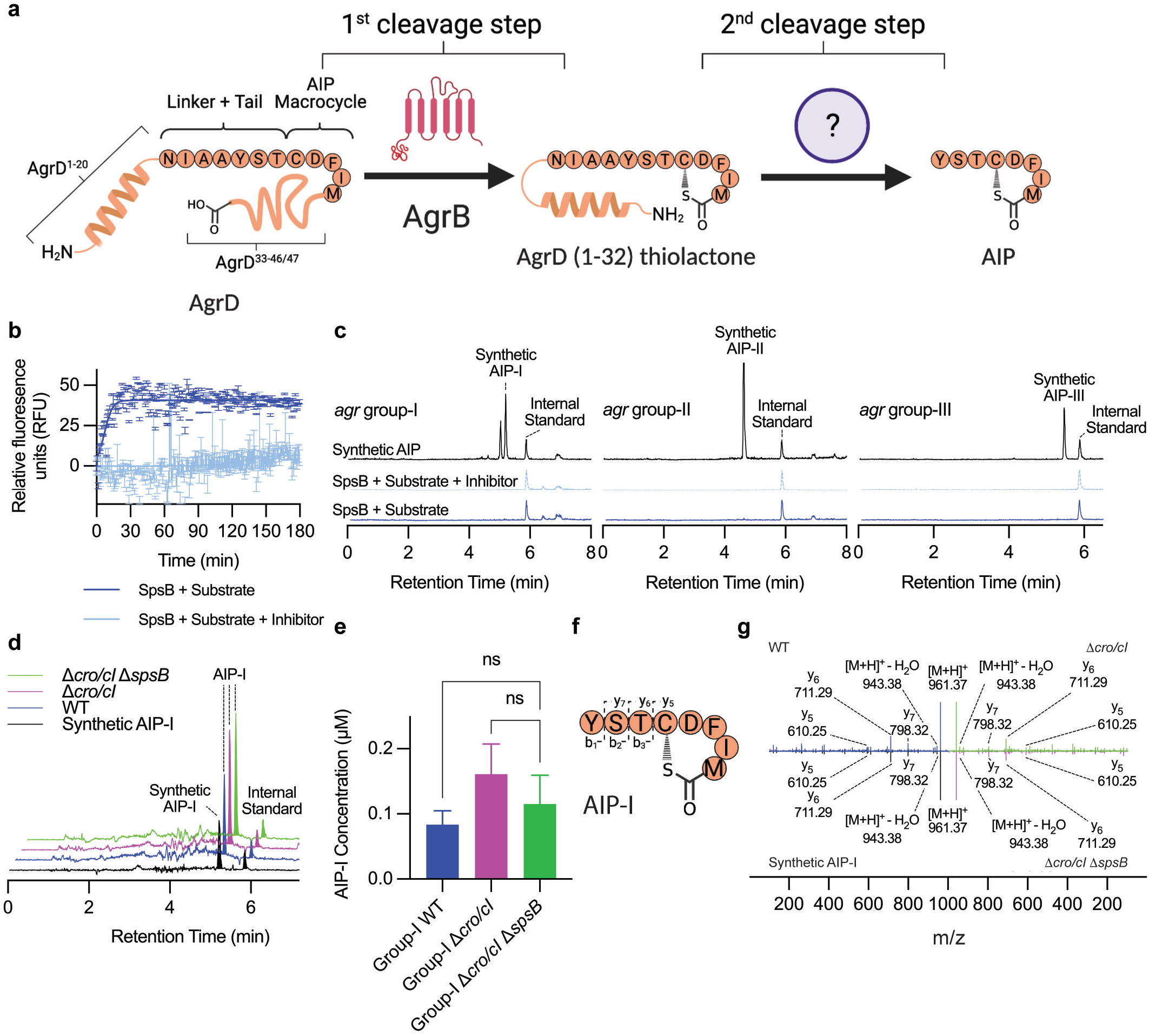
SpsB does not directly participate in AIP biosynthesis. **A**. Overview of the AIP biosynthetic pathway. AgrD (group-I sequence shown) is processed by AgrB to yield the biosynthetic intermediate, AgrD (1-32) thiolactone. This intermediate is converted into the mature AIP through the action of a second protease whose identity remains unclear. **B**. SpsB cleavage of a validated fluorogenic substate, DABCYL-SceD-EDANS, in the presence and absence of a known SpsB inhibitor, M131. SpsB proteoliposomes were incubated with the substrate ± inhibitor and the cleavage reaction monitoring by fluorescence (510 nm) over time. Data presented as the mean ± s.d. (n=3 replicates). **C**. In vitro assay for AIP production. SpsB proteoliposomes were treated with indicated AgrD (1-32) thiolactone intermediates in the presence or absence of M131 inhibitor. Reaction mixtures were then subjected to a solid phase extraction (SPE) step, prior to which an internal standard (AIP derivative) was added, and analyzed by LC-MS. Shown are the extracted ion currents (EIC) traces for the expected AIP and the internal standard. A synthetic AIP treated with empty liposomes served as a positive control (top trace). **D**. Cell based assay for AIP production. Indicated *S. aureus* strains (group-I background) were grown for 8h (the optimal timepoint for AIP-I production in the wild-type strain) at which point an internal standard was added and the media subjected to SPE and analyzed by LC-MS. Shown are the EIC traces for AIP-I and an internal standard. As a control, a synthetic AIP-I was added to media and subjected to the same purification protocol (black trace). **E**. AIP-I produced by indicated *S. aureus* strains was quantified by LC-MS using a standard curve approach employing synthetic AIP-I. Data presented as the mean ± s.d. (n=3-4 biological replicates). **F-G**. Comparative LC-MS/MS analysis of AIP-I produced by WT, Δ*cro/cI*, and Δcro/cI/Δ*spsB* variants of group-I *S. aureus* and a synthetic AIP-I standard. Color coding as in panel D.

Allelic variation within the *agr* operon, occurring as a result of hypervariability in *agrB, agrD*, and *agrC*, results in four distinct *agr* specificity groups within *S. aureus*.^19, 23, 24^ Each allelic variant produces a unique AIP-AgrC pairing. While cognate AIP/AgrC interactions are agonistic, non-cognate interactions are generally cross inhibitory to *agr* QS, with the exception of the closely related group-I and group-IV AIPs which cross activate.^19, 23^ Furthermore, biosynthesis of each unique AIP is performed through the proteolysis of a unique AgrD by a unique AgrB.^15^ While the size and hydrophobic nature of the AIP macrocycle are conserved among the specificity groups, AIPs derived from group-I/IV, group-II, and group-III are divergent in tail length and sequence.^19, 23, 24^ Structure-activity studies on AIPs reveal that shared structural motifs are necessary for AgrC binding, while divergent sequences determine if, upon binding, AIP acts as an agonist or an antagonist.^10, 25, 26^

Despite decades of research into the *agr* response, the AIP biosynthetic pathway has not been fully elucidated (**Figure 1A**). The second step of AIP maturation—cleavage of the N-terminal domain from the AgrD (1-32) thiolactone intermediate—is poorly understood relative to the rest of the system.^7^ Biochemical studies on model peptides derived on the group-I AgrD sequence indicate the signal peptidase I, SpsB, can perform this second cleavage step.^27^ While the involvement of SpsB in AIP maturation would be broadly consistent the enzyme’s known function in peptide secretion,^27, 28^ definitive *in vitro* and *in vivo* data establishing a role for this enzyme in AIP biosynthesis, including whether it is involved in the maturation of the group II-IV AIPs, is still lacking.

Recently, a putative integral membrane Abi-domain/M79 metalloprotease has been identified as a novel effector of *agr* QS in *S. aureus*.^29, 30^ Genetic studies indicate that this protein, designated MroQ (membrane protease regulator of *agr* QS), is required for *agr* activity and virulence production in a group-I *S. aureus* strain. These studies clearly implicate MroQ in the *agr-I* response, however, the precise function of the protein has not been definitively assigned. Indeed, Cosgriff *et al*^*29*^ propose a role for MroQ in AgrD processing and/or transport, while Marroquin *et al*^*30*^ conclude that MroQ is not directly involved in AgrD processing, but rather plays some other regulatory role.

In this study, we set out to characterize the second step in AIP maturation using a combination of biochemical and genetic approaches. Our studies argue against a major role for SpsB in AIP biosynthesis. By contrast, we provide evidence that MroQ catalyzes the second proteolytic cleavage step in the maturation of AIP-I/IV and AIP-II, but, surprisingly, not AIP-III. We also define the molecular features within the group-I and -II substrates that guide MroQ cleavage activity. As part of this effort, we successfully reconstitute much of the *agr* QS circuit using purified components, a first for a system of this type. Taken together, these results suggest a novel point of evolutionary divergence between the *S. aureus* specificity groups and underscore MroQ as a key component of the *agr* response.

## Results

### SpsB does not efficiently cleave AIP biosynthetic intermediates

Previous work has suggested a role for the membrane anchored protease, SpsB, in *agr* signaling.^27^ Notably, this study employed a truncated, soluble version of the enzyme and a short synthetic peptide substrate spanning the AgrD-I (1-32) thiolactone cleavage site. Thus, it remained to be determined whether the full-length, membrane-anchored enzyme can successfully process the native substrate, AgrD-I (1-32) thiolactone, into the mature AIP. It is also unclear if SpsB has any role to play in the maturation of AIPs from the other *S. aureus* specificity groups.

To address these questions, we overexpressed full-length *S. aureus* SpsB in *E. coli*. and then reconstituted the detergent solubilized purified protein into liposomes containing 1-palmitoyl-2-oleyl-phosphatidylcholine (POPC) and 1-palmitoyl-2-oleyl-phosphatidylglyerol (POPG) (**Figure S1A**,**B**). The bioactivity of these proteoliposomes was tested using a known synthetic SpsB substrate derived from the signal peptide sequence of *Staphylococcus epidermidis* pre-SceD, conjugated to an N- and C-terminal FRET pair DABCYL/EDANS (**Figure S1C**).^31^ Efficient cleavage of this peptide was observed in the presence of the SpsB proteoliposomes, activity that was abolished in the presence of the known SpsB inhibitor M131 (**Figures 1B and S1D**,**E**).^32^ We then incubated the SpsB-proteoliposomes with purified native AIP maturation substrate; AgrD (1-32) thiolactone from either *agr* group-I, group-II, or group-III *S. aureus* (**Figure S2**). AgrD-I (1-32) thiolactone and AgrD-III (1-32) thiolactone were generated recombinantly using an intein fusion strategy, whereas AgrD-II (1-32) thiolactone was chemically synthesized.^15^ In each case, the SceD substrate was also added as an internal control. Surprisingly, we did not observe the production of the native AIP in any of these reconstitution experiments, despite the fact that the SceD spike-in was successfully cleaved in each case (**Figure 1C, S3, S4A**).

Given this unexpected biochemical result, we were eager to determine whether deletion of *spsB* gene has any impact on AIP maturation in *S. aureus* cells. This experiment is complicated by the key role of SpsB plays in *S. aureus* viability.^33^ Fortunately, this dependency can be circumvented by de-repression of a putative ABC transporter, achieved through knockout of the transcriptional repressor *cro/cI*.^33^ Thus, we generated a mutant *S. aureus* group-I strain lacking *cro/cI* and *spsB* and tested for the production of AIP-I using a sensitive LC/MS approach for detecting the peptide secreted into the growth media. Consistent with the biochemical data, wild-type, Δ*cro/cI*, and Δ*cro/cI:*Δ*spsB* knockout strains exhibited similar levels of endogenous AIP-I production (**Figure 1D-G and S4B-D**). Together, these results argue against SpsB being the AIP maturation protease and point to the involvement of another membrane protease in the second biosynthetic step.

### MroQ activity is essential for AIP-I and AIP-II production, but not AIP-III

The recently identified *S. aureus* integral membrane protease, MroQ, presented itself as an attractive alternate candidate to SpsB in driving the second step in AIP maturation. This enzyme has been implicated in the group-I *agr* response, although the exact role it plays remains unclear.^29,30^ We decided to expand investigation of MroQ to include AIP biosynthesis in *S. aureus agr g*roups I, II, and III. Group-IV was not examined in this study, as *agrB, agrD*, and *agrC* from group-I and group-IV are highly conserved and AIP-I and AIP-IV, which differ by only a single amino acid, are cross-activating for AgrC-I and AgrC-IV, respectively.^23^ Thus, we can assume that AIP biosynthesis in group-I and group-IV operate via the same pathway and mechanism.

We began by performing a series of genetic complementation experiments analogous to those carried out in the initial studies implicating MroQ in the group-I *agr* response.^29, 30^ In the current study, we generated Δ*mroQ* knockout strains for *S. aureus agr* group-I, -II, and -III genetic backgrounds. A Cd^2+^ inducible expression plasmid,^34^ containing *mroQ*, an inactive *mroQ* mutant, or no insert, was then transduced back into the various Δ*mroQ* and wild type strains. We observed no major difference in growth rates for the three Δ*mroQ* knockout strains harboring these complementation plasmids relative to the corresponding WT strains (**Figure 2A**). This is consistent with the proposed role of *mroQ* in the *agr* circuit, as *agr* QS is non-essential for *S. aureus* viability.^35^

**Figure 2.**
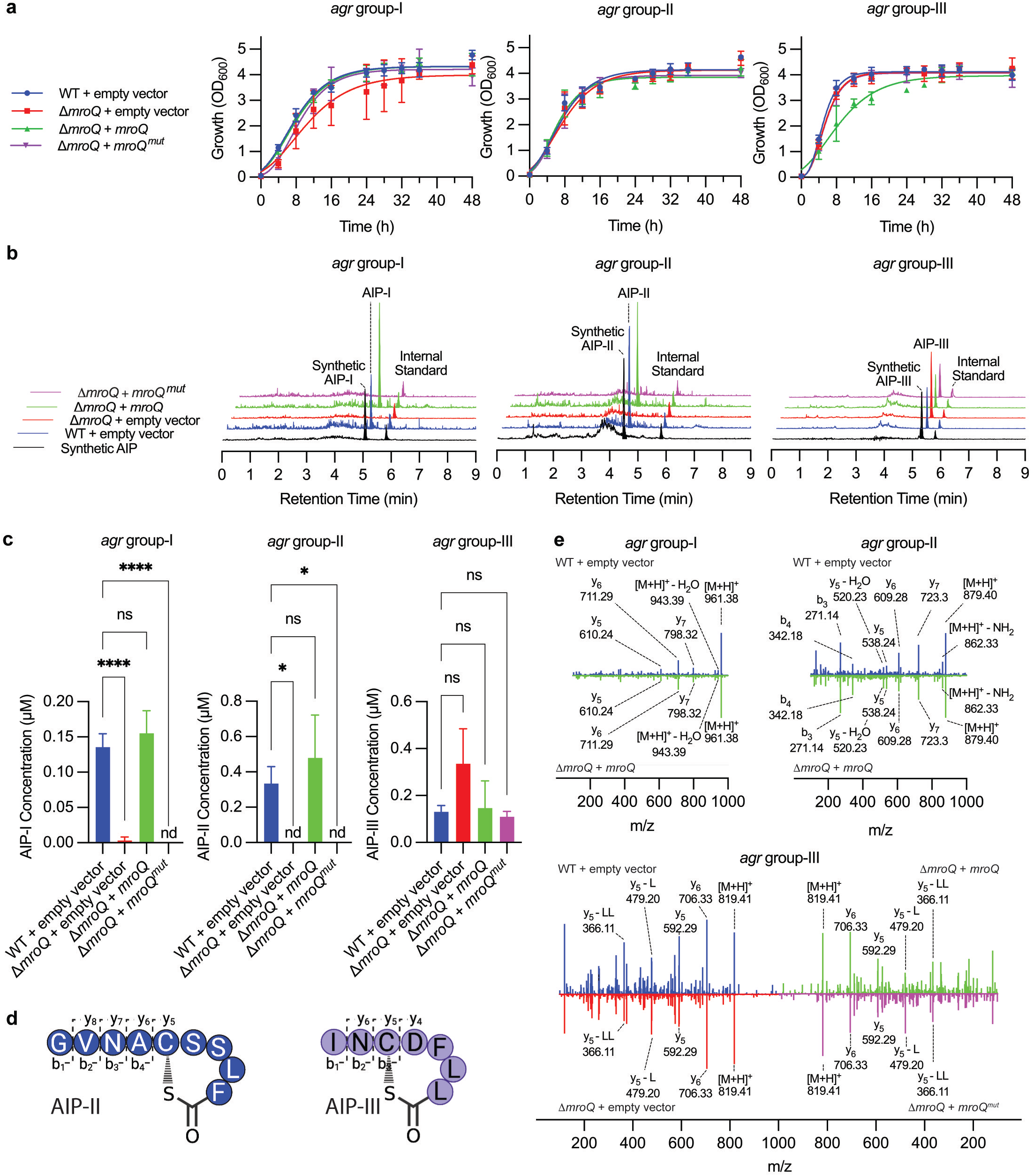
Genetic complementation approach to explore the role of *mroQ* in AIP biosynthesis. **A**. Relative growth of WT and Δ*mroQ* strains containing inducible plasmids expressing *mroQ, mroQ*^*mut*^, or an empty plasmid, generated in *agr* group-I, -II, and -III background strains over 48h, monitored by OD_600_. Data presented as the mean ± s.d. (n=3 biological replicates). **B**. Cell based assay for AIP production. Indicated *S. aureus* strains were grown for 8h (group-I) and 16h (group-II, group-III) at which point an internal standard was added and the media subjected to SPE and analyzed by LC-MS. Shown are the EIC traces for the AIPs and the internal standard. As a control, a synthetic AIP was added to media and subjected to the same purification protocol (black traces). **C**. AIP levels produced by indicated *S. aureus* strains were quantified by LC-MS using a standard curve approach employing synthetic AIPs. Data presented as the mean ± s.d. (n=3 biological replicates). **D-E**. Comparative LC-MS/MS analysis of AIPs generated by wild-type and mutant-complemented strains. Color coding as in panel C.

To assess the impact of *mroQ* deletion on AIP production, growth media was taken from the various *S. aureus* strains at periodic timepoints and the levels of AIP quantified by LC-MS using a standard curve approach (**Figure S5A**,**B**). Building on previous findings,^29, 30^ we found that deletion of the protease not only abolished AIP production in group-I strains relative to WT, but also profoundly reduced AIP levels in the group-II strain (**Figure 2B,C**). By contrast, deletion of *mroQ* had no significant effect on AIP production in the *S. aureus* group-III background. Use of a β-lactamase reporter cell assay also indicated that MroQ was needed for AIP production in groups -I and -II, but not group-III (**Figure S5C**). Importantly, complementation with wild-type *mroQ*, but not an inactive mutant,^29, 30^ rescued AIP production in *agr* group-I and group-II Δ*mroQ* knockout strains to near-WT levels (**Figure 2B-E**). Taken together, these studies extend the previous work in this area by showing that MroQ activity is required for both AIP-I and AIP-II production, whereas the group-III *agr* response seems to be independent of this enzyme.

### MroQ efficiently cleaves AgrD (1-32) thiolactones *in vitro* to yield mature AIPs

While the above genetic studies clearly link MroQ to AIP-I/II production, they do not reveal whether the protease is directly involved in the AgrD-I/II (1-32) thiolactone processing step, versus it playing an indirect role by regulating the activity of some other factor. To discriminate between these possibilities, we performed biochemical studies on the system using purified components. Full-length MroQ, as well as the inactive version of the enzyme MroQ^mut^,^29, 30^ were successfully over-expressed as MBP-fusions in *E. coli* (**Figure S6A**). Following purification, the detergent-solubilized proteins were then reconstituted into proteoliposomes using the same lipid system employed for SpsB. We then incubated these MroQ-proteoliposomes with the purified AgrD (1-32) thiolactones from groups I-III and analyzed the reaction mixtures using our quantitative LC-MS assay (**Figure 3A**). Efficient and specific cleavage of AgrD (1-32) thiolactone from *agr* groups I and II was observed in the presence of MroQ, generating AIP-I and AIP-II respectively as determined by LC-MS and MS/MS (**Figure 3B-D, Figure S6B**). Notably, MroQ^mut^ failed to catalyze substrate cleavage under equivalent conditions (**Figure S6B)**. Exposure of the group-III AgrD (1-32) thiolactone to the MroQ-proteoliposomes did not lead to efficient production of AIP-III, rather the major species observed in this reaction corresponded to a mis-cleavage product in which an additional Tyr is appended to the AIP-III N-terminus (**Figure 3B, Figure S6B-D**). These biochemical data demonstrate that MroQ is able to efficiently catalyze the second proteolytic cleavage step in AIP-I and AIP-II biosynthesis. By contrast, and in keeping our genetic studies, the biochemical data argue against a major role for this enzyme in the AIP-III maturation process.

**Figure 3.**
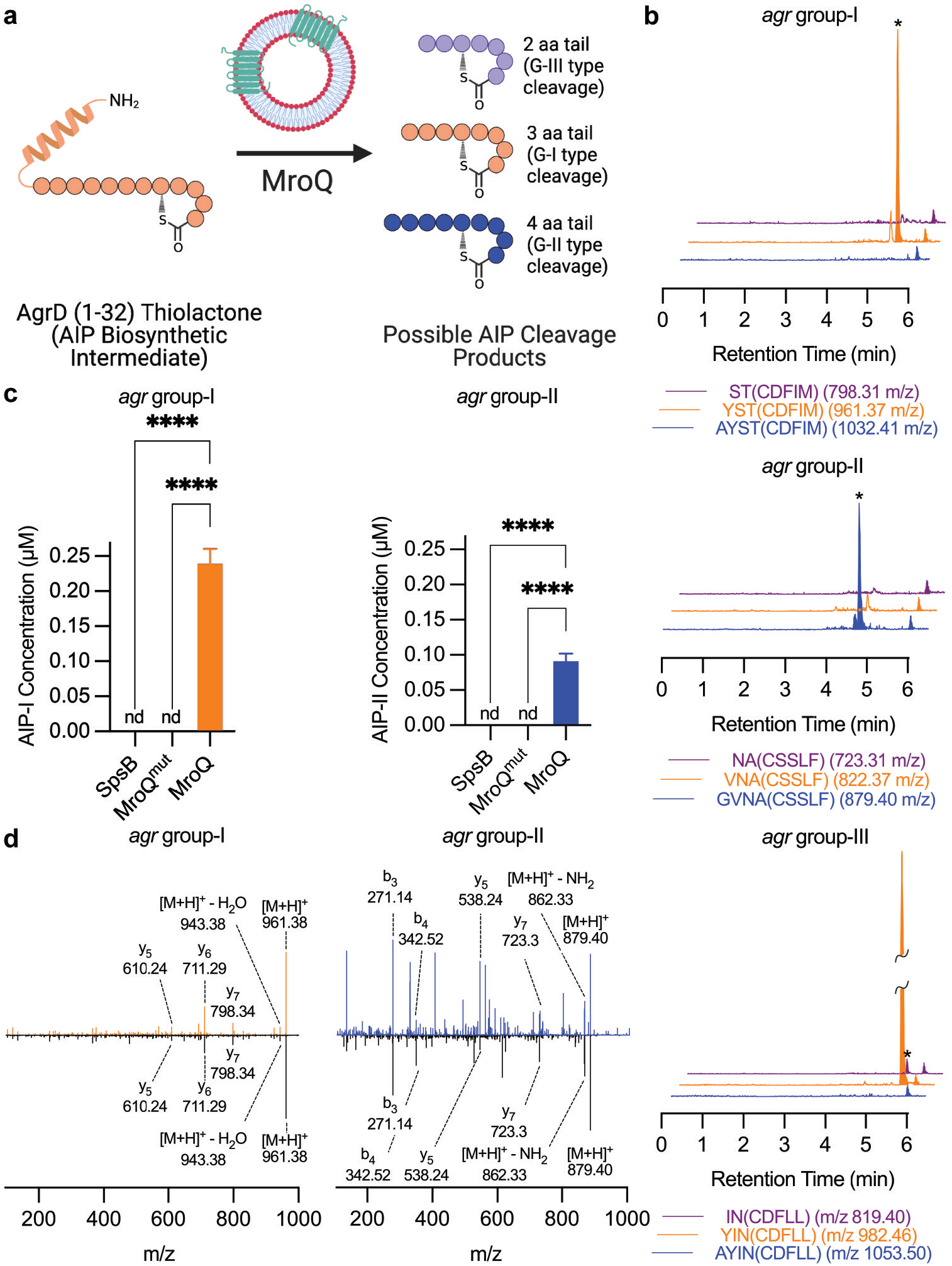
*In vitro* reconstitution of MroQ. **A**. Overview of *in vitro* cleavage assay assessing cleavage of AgrD (1-32) thiolactone substrates by MroQ reconstituted into proteoliposomes. **B**. MroQ proteoliposomes were combined with indicated AgrD (1-32) thiolactone intermediates. Reaction mixtures containing a spike-in internal standard were then subjected to a SPE step and analyzed by LC-MS. Shown are the EIC traces for the possible cleavage products and the internal standard. * Indicates correct AIP product from each specificity group. **C**. AIP-I and -II levels produced by MroQ, MroQ^mut^ or SpsB proteoliposomes were quantified by LC-MS using a standard curve approach employing synthetic AIPs. Data presented as the mean ± s.d. (n=3 biological replicates). **D**. Comparative LC-MS/MS analysis of AIPs produced from MroQ proteoliposome cleavage assay (top) and synthetic standards (bottom).

Next, we asked whether MroQ could work in conjugation with AgrB to produce mature AIP-I and AIP-II from the corresponding AgrD polypeptide precursors (**Figure 4A**). A reconstituted system was again employed, in this case combining recombinant AgrD peptides with MroQ-and AgrB-proteoliposomes, the latter prepared as previously described (**Figure S7A**).^15^ Employing our LC-MS readout, we observed the generation of mature AIP-I and AIP-II when the corresponding AgrD precursors were treated with the cognate AgrB protease in the presence of MroQ (**Figure 4B-E**). Thus, AgrB and MroQ can work in tandem to produce a mature AIP. The success of this experiment represents the first formal reconstitution of a complete AIP biosynthetic pathway.

**Figure 4.**
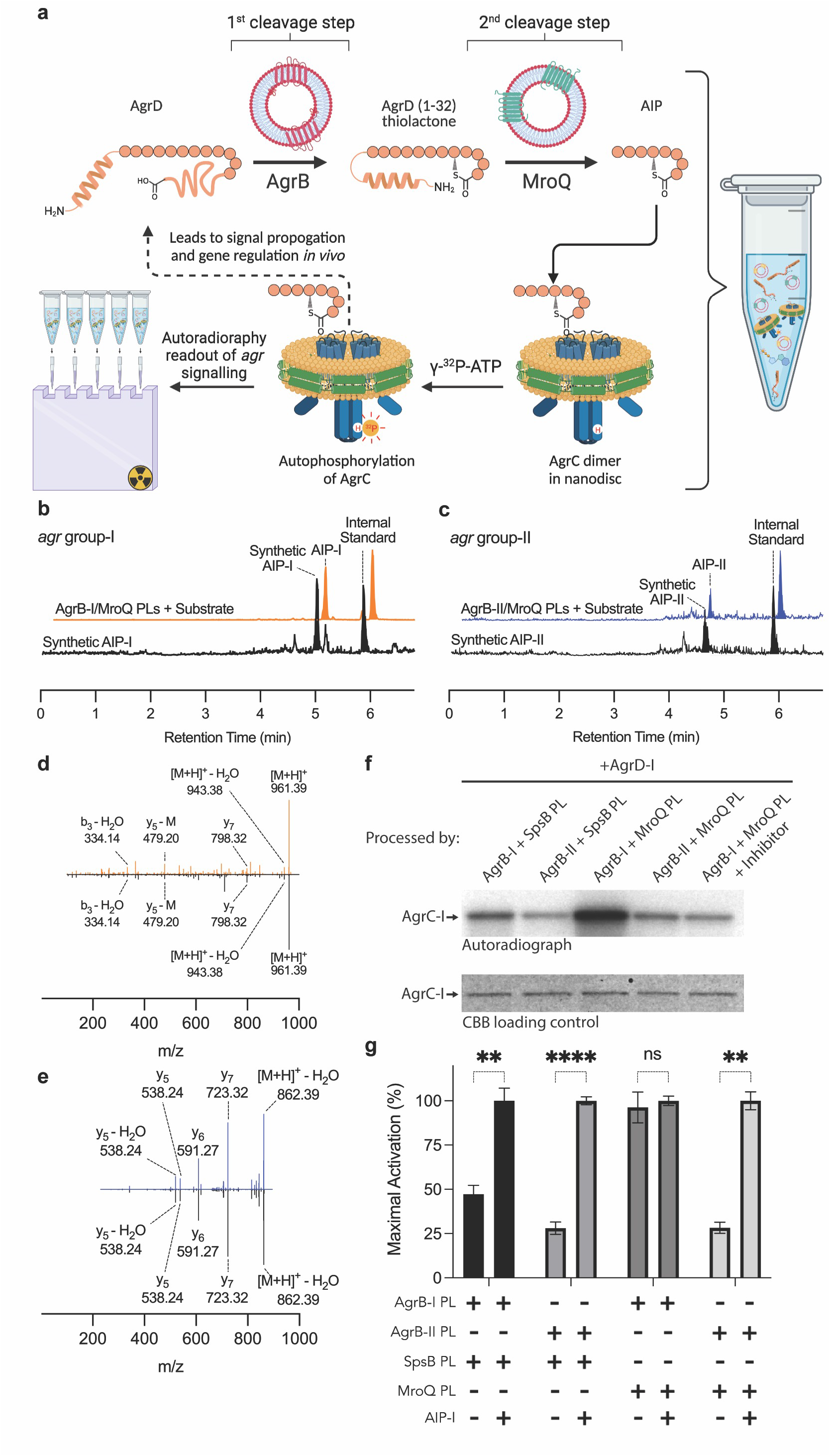
*In vitro* reconstitution of AIP biosynthesis and sensing. **A**. Overview of one-pot *in vitro* AIP biosynthesis and sensing assay. Input AgrD is processed by AgrB and MroQ proteoliposomes into mature AIP which then binds and activates nanodisc-reconstituted AgrC. **B-C**. Reconstituted AIP biosynthesis. AgrD-I or AgrD-II was incubated with cognate AgrB and MroQ proteoliposomes. Reaction mixtures containing a spike-in internal standard were then subjected to a SPE step and analyzed by LC-MS. Shown are the EIC traces for the expected AIP and the internal standard. A synthetic AIP treated with empty liposomes served as a positive control (bottom trace). **D-E**. LC-MS/MS analysis of AIP-I (D) and AIP-II (E) products of the *in vitro* biosynthesis reaction (blue) along with the corresponding synthetic standards (black). **F**. Autoradiography of AgrC-I autophosphorylation employing γ-^32^P-ATP. All samples contained AgrD and AgrC in addition to the indicated proteins. Inhibitor refers to a known tight binding antagonist of AgrC-AIP interaction. Note, AgrC-I is known have basal autokinase activity.^18^ **G**. Quantification of AgrC-I autokinase activity as determined by densitometry analysis of the autoradiographs. All samples contained AgrD and AgrC in addition to the indicated proteins and peptides. Maximal activation of AgrC-I was determined by addition of synthetic AIP-I to the mixtures. Data presented as the mean ± s.d. (n=3 biological replicates).

Encouraged by these reconstitution experiments, we decided to explore the feasibility of establishing a one-pot *in vitro* system comprising both the biosynthetic and AIP-sensing arms of the *agr* response. For this we employed our previously developed lipid nanodisc-embedded recombinant AgrC-I dimers,^14^ along with proteoliposomes containing the complete AIP biosynthetic pathway and precursor peptide (**Figure 4A & S7A**,**B**). Remarkably, in the presence of ATP-γ-^32^P, we observed robust AgrC-I autophosphorylation that was dependent upon the presence of all four protein components (**Figure 4F**,**G**). Moreover, AgrC-I activation was abolished in the presence of a known orthosteric inhibitor, QQ-3,^36^ confirming that the AIP produced *in situ* was engaging the cognate AgrC-I sensing pocket. To our knowledge, this represents the first full *in vitro* reconstitution of a bacterial quorum sensing pathway, recapitulating ribosomally synthesized and post-translationally modified peptide (RiPP) biosynthesis, receptor engagement, and receptor activation simultaneously under physiologically relevant conditions to achieve bioequivalent signal output in a single system.

### MroQ recognizes specific sequence motifs in AgrD-I/II-(1-32)-thiolactone

We next turned to elucidating the molecular features dictating MroQ cleavage specificity, which we note must occur at different positions in the AgrD-I/II (1-32) thiolactone substrates in order to generate AIPs of the correct length - mature AIP-I is an 8mer while AIP-II is one residue longer (**Figure S8**). Since the enzyme is fully conserved among the *S. aureus agr* specificity groups, we imagined that cleavage specificity must be driven by differences in the substrate. The AIP biosynthetic intermediate, the AgrD (1-32) thiolactone, contains three structural domains: the N-terminal amphipathic helix, the linker/AIP-tail region (which contains the MroQ cleavage site), and the AIP-macrocycle (**Figure S8**). The linker/AIP tail region is the most divergent between the different groups, making it the prime candidate as the specificity driver.

To test this possibility, we generated chimeric AgrD (1-32) thiolactone peptides containing the sequence of the linker domain from one specificity group, inserted into a peptide containing the N-terminal amphipathic helix and AIP-macrocycle sequences from a different specificity group (**Figure 5A & S9A**,**B**). These peptides were generated recombinantly by a similar intein fusion method used to generate native AgrD (1-32) thiolactones. For example, the AgrD-II-I-II (1-32) thiolactone chimera has the group-I linker sequence flanked by group-II sequences. These chimeras were then incubated with MroQ-proteoliposomes and the cleavage products analyzed by LC/MS and MS/MS (**Figure 5B-F**). The linker sequence from *agr* group-I directed MroQ to cleave the II-I-II chimeric intermediate at a position three amino acids from the AIP macrocycle yielding the chimeric AIP product, AIP-I-II, bearing a tail identical to AIP-I (**Figure 5B,D,F**). Similarly, the linker sequence from group-II intermediate directed cleavage of the chimeric substrate AgrD-I-II-I (1-32) thiolactone to yield AIP-II-I as the major product, indicating that the group-II linker sequence directs cleavage in AIP-II biosynthesis (**Figure 5C,E,F**).

**Figure 5.**
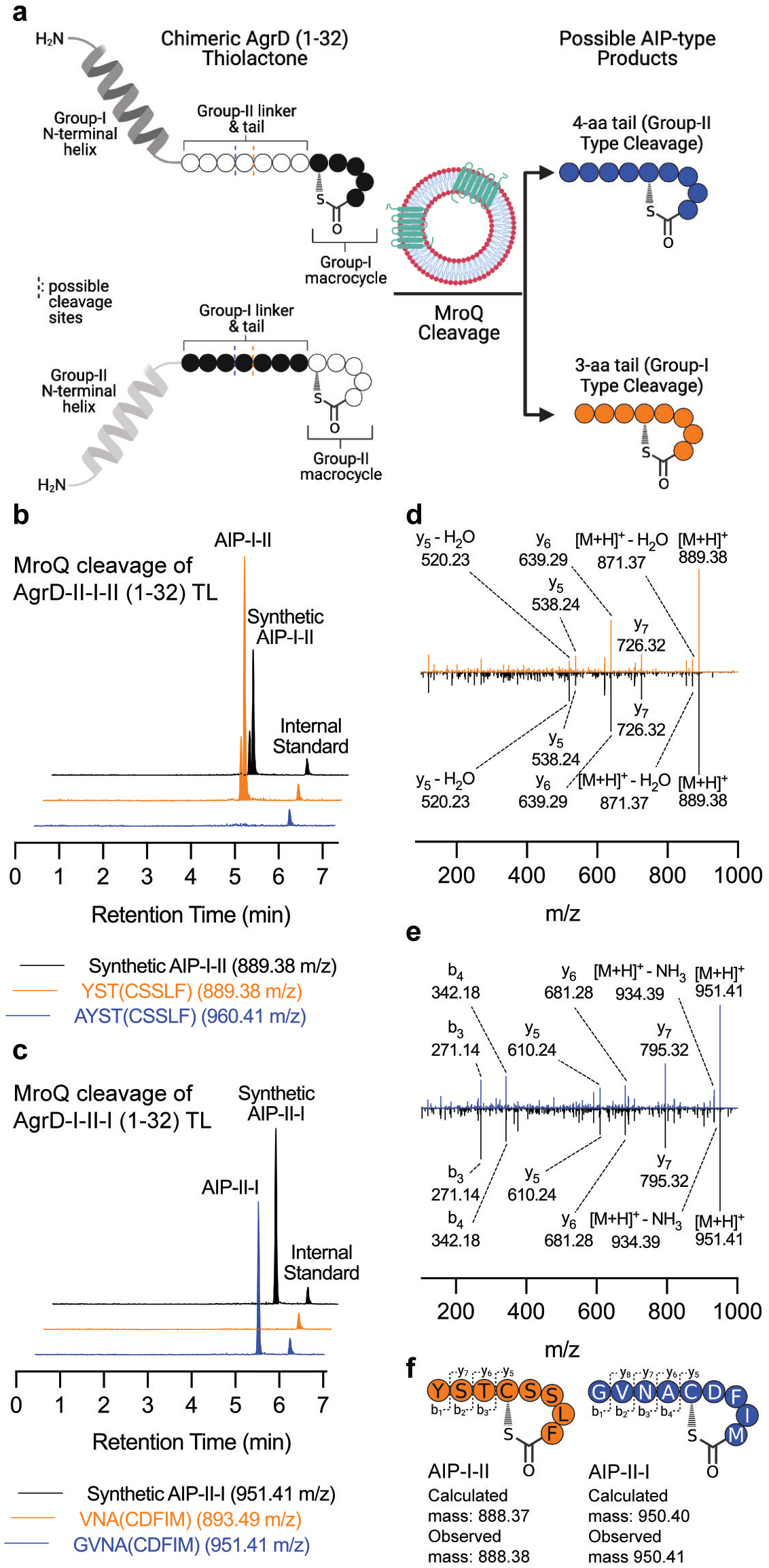
The AgrD-I/II linker domain dictates MroQ specificity. **A**. Overview of the in vitro MroQ cleavage assay employing chimeric AgrD-I/II substrates in which the linker-tail region of one specificity group is swapped for another. **B-C**. MroQ proteoliposomes were treated with indicated chimeric AgrD (1-32) thiolactone intermediates; AgrD-II-I-II (1-32) thiolactone (panel B) and AgrD-I-II-I (1-32) thiolactone (panel C). Reaction mixtures containing a spike-in internal standard were then subjected to a SPE step and analyzed by LC-MS. Shown are the EIC traces for the possible AIP cleavage products and the internal standard. Synthetic chimeric AIPs treated with empty liposomes served as positive controls (black traces). **D-F**. LC-MS/MS analysis of AIP-I-II (D, orange) and AIP-II-I (E, blue) products along with the corresponding synthetic standards (black).

To validate these biochemical findings, we generated *S. aureus* strains harboring equivalent chimeric *agrD* mutants. The desired chimeric *agrD* sequences were introduced into the *agr* null strain, RN7206, along with cognate *agrB, agrC and agrA* sequences, paired based on the identity of the *agrD* AIP-macrocycle domain (**Figure S10**). Importantly, the *agrC* sequences used in these experiments harbored a mutation (R238K) that leads to constitutive AgrC activity,^37^ ensuring continuous transcription of the mutant RNAII regardless of the agonist or antagonist activity of secreted chimeric AIP analogs (**Figure 6A**). Consistent with the *in vitro* studies employing chimeric substrates, the identity of the linker region was observed to direct cleavage site position. Thus, expression of AgrD-II-I-II led to the secretion of the chimeric AIP-I-II, with an AIP-II macrocycle and a three amino acid AIP-I tail consistent with group-I type cleavage (**Figure 6B,D & 5F**). Similarly, expression of AgrD-I-II-I led to the secretion of the chimeric AIP-II-I, with a four amino acid AIP-II tail consistent with group-II type cleavage (**Figure 6C,E & 5F**). Based on these data, we conclude that the AgrD linker motif sequence dictates specificity of MroQ mediated cleavage in AIP-I/II biosynthesis.

**Figure 6.**
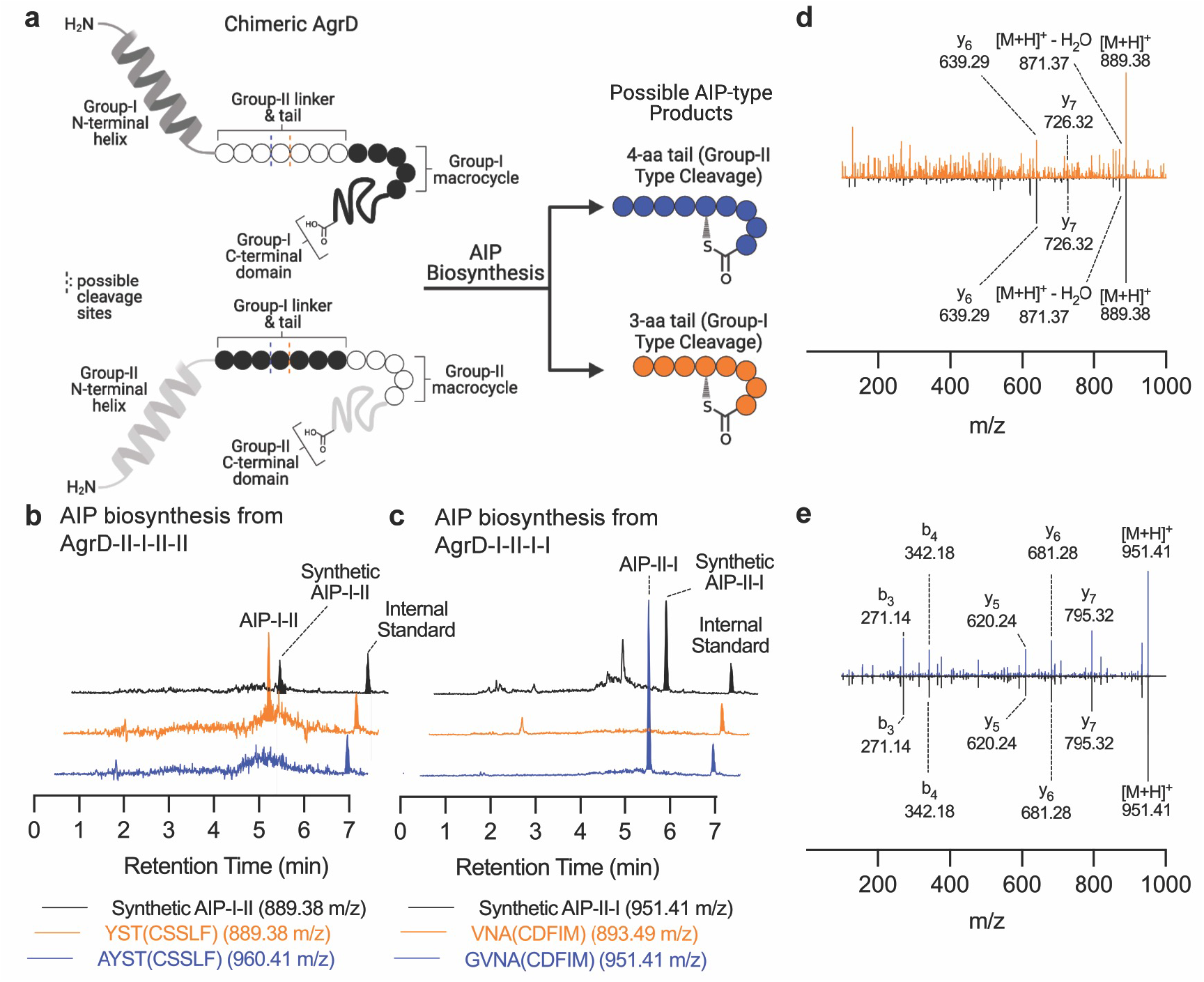
The AgrD-I/II linker domain directs AIP biosynthesis in *S. aureus* cells. **A**. Chimeric AgrD peptides containing the N-terminal helix domain, AIP macrocycle domain, and C-terminal domain from one *agr* specificity group, and a linker domain sequence from a different specificity group, are expressed under P2 promoter control in an *agr*-null *S. aureus* background. Identity of secreted chimeric AIPs determined by LC-MS. **B-C**. EIC traces of chimeric AIP cleavage products and internal standard from LC-MS analysis of growth media from *S. aureus cells* expressing AgrD-II-I-II-II (panel B) and AgrD-I-II-I-I (panel C) chimeras. Synthetic chimeric AIPs served as positive controls (black traces). **D-E**. LC-MS/MS analysis of AIP-I-II (D, orange) and AIP-II-I (E, blue) products isolated from growth media along with the corresponding synthetic standards (black).

## Discussion

The *S. aureus agr* quorum sensing circuit plays a central role in the regulation of virulence in this human pathogen, and as a consequence has been the subject of intense study for decades.^2, 7^ Despite this, our understanding of the biosynthetic pathway that leads to the generation of the secreted AIP signaling molecules has remained incomplete. In this study, we employed biochemical reconstitution and genetic complementation strategies to show that the integral membrane protease, MroQ, plays a direct role in the biosynthesis of the AIPs from *S. aureus agr* specificity groups I and II. Our data indicate that this enzyme can perform the second step in the maturation process, namely cleavage of the AgrD-I/II (1-32) thiolactone intermediates, and that the specificity of this processing step is dictated by the linker region in the substrates. Given the high sequence homology between the group-I and group-IV *agr* systems, we propose that MroQ also plays a direct role in the maturation of the latter. By contrast, we find no evidence that MroQ is directly involved the biosynthesis of the group-III AIP. Our biochemical and genetic data also argue against the involvement of the signaling peptidase, SpsB, an enzyme previously implicated in AIP-I biosynthesis. Collectively, these data shed new light on the mechanism of AIP biosynthesis and reveal an unexpected divergence in this maturation process between the *agr-*III system and the other specificity groups.

Several recent reports have highlighted the involvement of MroQ in *S. aureus* virulence. In a key pair of studies, Cosgriff *et al*. and Marroquin *et al*. used genetic approaches to show that MroQ is required for *agr* activity and virulence production in a group-I *S. aureus* strain.^29, 30^ However, these investigators stopped short of assigning a specific biochemical role for the protein, leaving it unclear whether the protease is directly involved in AIP maturation, or plays some other function in the *agr* signaling circuit, for example by modulating AIP sensing. Adding to the uncertainty, a recent study has even suggested that MroQ might hydrolyze the AIP, leading to the downstream inactivation of AgrC and by extension *agr* QS.^38^ Our biochemical studies clarify this picture by firmly establishing a direct role for MroQ in the second step of AIP biosynthesis, at least for *agr* groups I and II. Moreover, we see no evidence that the enzyme metabolizes the mature AIP.

We also investigated whether the signal peptidase, SpsB, could convert the AIP(1-32)-thiolactone intermediates from groups I-III into the corresponding AIPs. While we were able to reconstitute active full-length enzyme, as evidenced by robust cleavage of a known substrate, our efforts to generate AIPs using SpsB-proteoliposomes were unsuccessful. This result stands in contrast to the work of Kavanaugh *et al*. who demonstrated that a soluble N-terminally truncated version of SpsB cleaves peptide mimics of the AgrD-I linker domain.^27^ Conceivably, the full-length, membrane-integrated protein might require some additional factor, not present in our reconstituted system, in order to cleave the membrane-associated AgrD (1-32)-thiolactone. However, the fact that we observed cleavage of a spike-in control substrate in these experiments shows that the enzyme can at least access substrates in this proteoliposomal context. Furthermore, our use of a genetic rescue system that allows deletion of *spsB* in group-I *S. aureus* cells, also failed to provide evidence supporting a role for this enzyme in AIP production. Taken together, our data suggest the SpsB is not required for AIP maturation.

While our biochemical and genetic experiments converge on a direct role for MroQ in AIP maturation in *S. aureus* groups I and II, they also argue against the involvement of this enzyme in the biosynthesis of AIP-III. Thus, we imagine that another, as yet unidentified, protease must be involved in the group-III system, i.e. it is divergent from the other specificity groups at the level of the biosynthetic pathway. Clues as to the origins of this divergence emerge from our structure-activity studies indicating that MroQ cleavage specificity is dictated by the linker region connecting the N-terminal amphipathic helical domain and the macrocyclic domain. In the case of the group-I and -II systems, sequence variation within this region appears sufficient to direct MroQ cleavage to the correct site, resulting in AIPs of different length. This relationship appears not to hold true for the MroQ:AgrD-III pairing. Conceivably, the sequence of the linker region in AgrD-III is incompatible with efficient and specific MroQ activity, for example it contains an arginine residue in place of the valine or isoleucine found in the other three groups. We also note that the N-terminal helical domain of AgrD-III differs at multiple positions from the other three groups (**Figure S8**). In particular, it contains substitutions that alter amino acid charge which might affect how this domain associates with the bacterial membrane. It is possible that either, of both, of these unique sequence characteristics direct AgrD-III (1-32) thiolactone intermediate to a different processing pathway.

Finally, our studies do not reveal where MroQ-mediated cleavage of the AgrD-I/II (1-32) thiolactone intermediates occurs relative to the plasma membrane. This has important implications for how the mature AIPs are secreted out of the cell. One can certainly imagine a scenario where the substrate enters the protease active site from the cytoplasmic side of the membrane, meaning that it is the mature AIP that subsequently crosses the membrane via a passive or active transport process. With respect to this, we note that MroQ does not share homology with known ABC transporters, however we cannot rule out that its proteolytic activity is somehow coupled to transport, bearing in mind that this is an integral membrane protein. Conversely, the substrate could enter the MroQ active site from the extracellular side. In this case, cleavage activity would release the mature AIP directly into the extracellular milieu. However, such a mechanism requires that the AgrD (1-32) thiolactone intermediate, or at least the linker region within in, somehow localizes to the extracellular side of the membrane in order to gain access to the enzyme active site. Again, both passive and active transport processes are possible.^7^ Ultimately, additional biochemical and likely structural studies will be required to fully address these outstanding questions. Regardless, our data reveal MroQ as a bona fide player in AIP biosynthesis and, moreover, suggest a novel point of evolutionary divergence between the *S. aureus* specificity groups.

## Acknowledgements

We thank current members of the Novick and Muir laboratories for discussions and comments. We thank Connie Wang for assistance in development and synthesis of substrates and mutant *S. aureus* strains. We thank Dr. John Eng and Dr. Venu Vandavasi from the Princeton Chemistry Mass Spectrometry Facility, and Dr. Saw Kyin from the Princeton Molecular Biology Proteomics Facility for their assistance. We thank Merck for the gift of M131. Figures created with Biorender. This work was supported by National Institutes of Health (NIH) Grant AI042783 to T.W.M.

## Author contributions

S.P.B., A.Z., and Q.X. performed all experiments. S.P.B., A.Z., Q.X., B.W., G.R., and R.N. assisted in substrate and strain generation. S.P.B., A.Z., Q.X., and T.W.M. analyzed all data. S.P.B, A.Z, and T.W.M. wrote the manuscript.

## Methods

### Materials and General Methods

All common buffering salts, guanidine hydrochloride (GdnHCl), Tris(2-carboxyethyl)phosphine hydrochloride (TCEP), phenylmethylsulfonyl fluoride (PMSF), sodium 2-sulfanylethanesulfonate (MESNa), isopropyl-β-D-thiogalactopyranoside (IPTG), Luria-Bertani (LB) broth, LB agar, Coomassie brilliant blue, Remel tryptic soy agar, yeast extract, and HisPur Nickel-nitrilotriacetic acid (Ni-NTA) resin were purchased from ThermoFisher Scientific (Waltham, MA). Tryptic soy broth was purchased from Fluka Analytical (Charlotte, NC). Casamino acids were purchased from Difco Biosciences, Inc. (San Diego, CA). β-Glycerophosphate was purchased from EMD Millipore (Billerica, MA). Trifluoroacetic acid (TFA) was purchased from Halocarbon (North Augusta, SC). Talon cobalt resin was from Clontech (Mountain View, CA). All lipids were purchased from Avanti Polar lipids (Alabaster, AL) and all detergents were obtained from Anatrace (Maumee, OH). EDANS-NovaTag resin was purchased from Novabiochem (Läufelfingen, Switzerland). Trityl-ChemMatrix resin was purchased from Biotage (Charlotte, NC). PyBOP was purchased from Oakwood Chemical (Estill, SC). HOBt was purchased from AnaSpec (Fremont, CA). Fmoc-amino acids were purchased from Matrix Innovation (Quebec, Canada). ATP-γ-^32^P was purchased from PerkinElmer (Waltham, MA). Oligonucleotides were purchased from Integrated DNA Technologies (Coralville, IA). Codon-optimized gene fragments were purchased from GENEWIZ (South Plainfield, NJ). Gibson Assembly Master Mix and all empty plasmids were purchased from New England Biolabs (Ipswich, MA). DH5α, BL21(DE3), and C43(DE3) *E. coli* strains were purchased from Invitrogen (Carlsbad, CA) and subcultured to develop stocks of competent cells. Ammonium persulfate (APS), 30% Acrylamide/Bis Solution 29:1, tetramethylethylenediamine (TEMED), criterion empty gel cassettes, and Bio-Beads SM2 were purchased from Bio-Rad (Hercules, CA). Sep-PaK Vac 1cc (50 mg) C18 cartridges were purchased from Waters (Milford, MA). Amicon Ultra-15 30 kDa MWCO centrifugal filter unit concentrators were obtained from MilliporeSigma (Burlington, MA). Chromatography columns were purchased from Environmental Express (Charleston, SC). DNA purification kits were purchased from Qiagen (Valencia, CA). All other reagents and solvents were purchased from Sigma-Aldrich (St. Louis, MO). All reagents and solvents were used without further purification.

The identity and composition of all plasmids and all *S. aureus* strains used in this study were validated by Sanger sequencing performed by GENEWIZ.

Cell disruption was performed using an ATA Scientific Instruments Avestin Emulsiflex C3 French Press (Caringbach NSW, Australia) or an Ultrasonic Liquid Processor Sonic Dismembrator (ThermoFisher). All SEC-FPLC described in this study was performed on a GE Healthcare AKTA PV-908 FPLC system (Chicago, IL) on Superdex S200 10/300 GL (Sigma-Aldrich) and Superose 6 10/300 GL (Sigma-Aldrich) columns. SDS-PAGE gels and autoradiographs were imaged on a LI-COR Odyssey Photoimager (Lincoln, NE) or a Cytiva Amersham ImageQuant 800 (Malborough, MA). Large scale peptide purification was performed by preparative-scale reverse-phase HPLC using a 2535 Quartenary Gradient Module (Waters) employing an XBridge Peptide BEH C18 OBD Prep Column, 300Å (10 μm, 19 × 250 mm) (Waters) at a flow rate of 20 mL/min. All other synthetic or expressed peptide substrates were purified and analyzed at small scale utilizing analytical-scale reverse-phase HPLC on an Agilent 1100 series HPLC system (Santa Clara, CA) employing an Eclipse Plus C18 column (5 μm, 4 × 150 mm) (Agilent) at a flow rate of 1 mL/min. For all reverse-phase HPLC purification or analysis, 0.1% TFA in water (HPLC solvent A) and 90% acetonitrile, 10% water, and 0.1% TFA (HPLC solvent B) were used as the mobile phases. Lyophilization of samples after HPLC or SPE was performed on a Millrock Technology MD85 lyophilizer (Kingston, NY).

Mass validation of peptides was performed by electrospray ionization mass spectrometry (ESI-MS). Proteins were characterized by SDS-PAGE, matrix-assisted laser desorption/ionization (MALDI) MS, and by LC-MS/MS following tryptic digestion. ESI-MS was performed on a 6120 Quadrupole LC/MS (Agilent) employing a 1260 Infinity LC system (Agilent) with a Zorbax 300SB-C18 column (3.5 μm, 4.6 × 100 mm) (Agilent) at a flow rate of 0.5 mL/min. MALDI was performed on a Bruker Daltonics ultrafleXtreme MALDI-TOF/TOF (Billerica, MA). Tryptic digests of SDS-PAGE resolved proteins were analyzed on a LTQ XL linear ion trap mass spectrometer (ThermoFisher) connected to an Easy nLC HPLC (ThermoFisher). Mass validation of peptide products from *in vitro* cleavage assays or *in vivo* endogenous production was performed by LC-MS/MS on a 6546 LC/Q-TOF (Agilent) employing a 1290 Infinity II LC system using an Atlantis dC_18_ column (5 μm, 4.6 × 100 mm) (Waters) at a flow rate of 0.75 mL/min. For this instrument, 0.1% formic acid in water was used as Solvent A, and 99.9% acetonitrile and 0.1% TFA was used as Solvent B. LC-MS/MS data was processed using the Masshunter Workstation (Agilent). Analysis of FRET-based cleavage assays was performed using a Molecular Devices SpectraMax M3 microplate reader (San Jose, CA). Analysis of nitrocefin cleavage in β-lactamase *agr* activation reporter cell assays was performed using a Multiskan GO 96-well plate reader and shaker incubator (ThermoFisher).

### Bacterial strains and plasmids

The bacterial strains and plasmids used in this study are listed in Supplementary Table 1. Primers used in this study are listed in Supplementary Table 2.

Knockout of *cro/cI*, s*psB, and mroQ* were performed using an in-frame deletion strategy in *agr* group-I (for all three genes), -II (*mroQ*), and -III (*mroQ*) *S. aureus* strain backgrounds, leveraging the pMAD shuttle vector double-crossover method as described in Arnaud *et al*. 2004.^39^ Knockouts were confirmed by PCR and sequencing (Genewiz). Complementation plasmids were generated by Gibson assembly using the inducible vector pCN51 and *mroQ* amplified from *S. aureus* gDNA, or catalytically impaired *mroQ*, with E141A and E142A mutations, obtained as a synthetic gene fragment (Genewiz). MroQ is fully conserved among the four *S. aureus agr* specificity groups, obviating the need for specificity-group specific MroQ expression. Plasmids were transformed into *E. coli* DH5α for plasmid amplification, purification (Qiagen), and sequence validation (Genewiz). Validated plasmids were transformed into *S. aureus* RN4220 by electroporation. Positive transformants were grown and lysed by staphylococcal phage 80α to generate a phage lysate for transduction into host *S. aureus* strains as outlined in Charpentier *et al*. 2004.^34^ All *S. aureus* strains containing transduced plasmids were generated by this method.

*S. aureus* mutants expressing chimeric AgrD peptides were generated by a chromosomal integration strategy outlined in Chen *et al*. 2014 in an *agr*-null background strain, RN7206.^20, 40^ Sequences containing *agrP2*-*agrB-I-agrD-I-II-I-agrC-I*^*R238K*^-*agrA*, or *agrP2*-*agrB-II-agrD-II-I-II-agrC-I*^*R238K*^-*agrA*, were generated and inserted into the SaPI1 integration suicide vector pJC1111 by overlap extension PCR. Plasmids were transduced into *S. aureus* via standard phage 80α transduction protocols outlined above.^34^ Positive chromosomal integrants at the SaPI1 chromosomal attachment (*attC*) site were selected with cadmium.

Cloning for expression of recombinant proteins was performed in *E. coli* DH5α, and expression of recombinant proteins was performed in *E. coli* BL21(DE3) and C43(DE3). *E. coli* codon optimized *His*_*5*_*-TEV-spsB* (*spsB*) and *MBP-mroQ-His*_*6*_ (*mroQ*) were obtained as codon-optimized synthetic DNA fragments (Genewiz). *His*_*6*_*-MBP-mroQ*^*E141AE142A*^*-HA* (*mroQ*^*mut*^) was generated by site-directed mutagenesis (NEB). *spsB, mroQ* and *mroQ*^*mut*^ sequences were PCR amplified and cloned into the NdeI and XhoI sites of a pET15b (*spsB*) and pET24b (*mroQ, mroQ*^*mut*^) vectors by Gibson assembly (NEB). Plasmids containing *agrB-I-His*_*6*_, *agrB-II-His*_*6*_, *agrC-I-His*_*6*_, *agrD-I-gyrA-His*_*7*_, and *agrD-II-gyrA-His*_*7*_ were previously generated.^14, 15^ Plasmids encoding *agrD-I(1-32)-gyrA-His*_*7*_ and *agrD-III(1-32)-gyrA-His*_*7*_ thiolactone were cloned between the NdeI and SapI sites of a modified pTXBI (IMPACT) vector (NEB), containing a C-terminal *Mxe* GyrA-His_7,_ via cloning protocols described in Wang *et al*. 2015 for AgrD-I and AgrD-II.^15^ Cloning, expression, and purification of MSP1D1 was performed via protocols described in Wang *et al*. 2014 for MSP1E3D1.^14^

### Expression and Purification of Recombinant MroQ, MroQ^mut^, SpsB, AgrB, AgrD, AgrC, and MSP1D1 Constructs

*MroQ*: MBP-MroQ, and MBP-MroQ^E141AE142A^ (MBP-MroQ^mut^) fusion constructs were expressed with MBP solubility tags and His_6_ purification handles in C43(DE3) *E. coli*. Note, the protein was found to be poorly behaved in the absence of the MBP tag. Bacteria were grown at 37 °C in four liters of LB medium with 50 μg/mL kanamycin to an optical density of 0.8 at 600 nm (OD_600_). Bacterial cultures were then cooled to 18 °C and expression of MBP-MroQ or MBP-MroQ^mut^ was induced by the addition of 0.5 mM IPTG. Protein was expressed overnight at 18 °C. Bacteria were harvested by centrifugation at 4,000 x g for 30 min. Bacterial pellets were resuspended in 10 mL of TBS lysis buffer / L LB media (20 mM Tris, 100 mM NaCl, 1mM PMSF, at pH 7.5). Bacteria was lysed by French press homogenization using a Avestin Emulsiflex C3 (ATA Scientific Instruments) through four passages of homogenized bacteria. Cell-wall debris was separated by centrifugation at 15,000 x g for 30 min. Membrane vesicles in the supernatant were separated from cytosolic proteins and materials by ultracentrifugation at 200,000 x g for 2 hours. Pelleted membrane fractions were resuspended in 5 mL of buffer (20 mM Tris, 100 mM NaCl, 2 mM TCEP, and 2% (w/v) n-Dodecylphosphocholine (Fos-Choline-12), at pH 7.5) containing detergent to solubilize membrane proteins. The resuspended membrane fraction was incubated overnight at 4 °C on a rotator. Solubilized membrane proteins were separated from membrane fragments by ultracentrifugation at 100,000 x g for 30 min. The supernatant was loaded on to Ni-NTA resin at a ratio of 1.5 mL resin bed / L expression and incubated for two hours at 4 °C on a rotator. Protein-bound resin was repacked in 25-mL Bio-Rad disposable plastic columns. Flow-through containing unbound protein was discarded and the column was washed with 15 column volumes (CV) of wash buffer I (20 mM Tris, 500 mM NaCl, 20 mM Imidazole, 1 mM TCEP, 0.05% (w/v) Fos-Choline-14 at pH 7.5) and 15 CV of wash buffer II (20 mM Tris, 100 mM NaCl, 30 mM Imidazole, 1 mM TCEP, 0.05% (w/v) Fos-Choline-14 at pH 7.5). Bound protein was eluted with 2 washes of 3 CV of elution buffer (20 mM Tris, 100 mM NaCl, 300 mM Imidazole, 1 mM TCEP, 0.05% (w/v) Fos-Choline-14 at pH 7.5). The elution was concentrated to 2 mL and further purified on a Superdex 200 10/300 GL size-exclusion chromatography column with 1.6 CV of running buffer (20 mM Tris, 100 mM NaCl, 1 mM TCEP, 0.05% (w/v) Fos-Choline-14, at pH 7.5). Peak fractions containing MBP-MroQ or MBP-MroQ^mut^ were pooled and concentrated, diluted to 10% glycerol, and snap frozen in N_2_ for storage at -80 °C. Analysis and mass validation of MBP-MroQ or MBP-MroQ^mut^ was performed by SDS-PAGE and tryptic digest LC-MS/MS (data not shown). *SpsB*: SpsB was expressed as a N-terminal His_5_ tagged recombinant protein. SpsB expression strain (BL21(DE2) *E. coli*) was grown at 37 °C in LB supplemented with ampicillin to OD_600_ = 0.5. Expression of SpsB was induced by addition of 1 mM IPTG. Bacteria was collected by centrifugation 3 hours post-induction and cell pellets were snap frozen. Upon thawing, cells were resuspended in 20 mL of lysis buffer (50 mM phosphate, 300 mM NaCl, 10 mM imidazole, 1 mM PMSF, pH 7.6). Note, addition of PMSF greatly improves the final purity of SpsB while having no effect on activity (data not shown). Bacteria were lysed by French press homogenizer and total protein was separated from cell-wall debris by centrifugation at 40,000 x g for 30 minutes at 4°C. Membrane vesicles containing SpsB were isolated by ultracentrifugation at 100,000 x g for 1 hour at 4°C. Membrane pellet was resuspended in lysis buffer, supplemented with 0.05% (w/v) DDM to solubilize membrane proteins, and incubated overnight at 4 °C. Soluble membrane proteins were separated from membrane debris by centrifugation at 50,000 x g for 30 min at 4°C. Total membrane protein fraction was incubated with TALON Co-NTA affinity resin for 2 hours at 4°C and washed with 5 CV of wash buffer I (lysis buffer, 0.05% DDM) and 5 CV of wash buffer II (lysis buffer with 30 mM imidazole, 0.05% DDM). The protein was then eluted with 2 CV elution buffer I (lysis buffer with 90 mM imidazole, 0.05%DDM) and 2 CV elution buffer II (lysis buffer with 300 mM imidazole, 0.05% DDM). Elutions were concentrated in 10 kDa MWCO concentrators and purified by size exclusion chromatography (SEC) on a S200 10/300 column in SpsB FPLC buffer (50 mM phosphate, 150 mM NaCl, 0.05% DDM, pH 7.5) over 1.6 CV. FPLC eluent was analyzed by SDS-PAGE. SpsB-containing fractions were concentrated in 10 kDa MWCO concentrators to 40 μM and supplemented with 10% glycerol. Aliquots were snap frozen and stored at -80 °C. Final SpsB products analyzed by SDS-PAGE and MALDI-ToF (data not shown). *AgrB, AgrC, and MSP1D1*: AgrB-I and AgrB-II were expressed as C-terminal His_6_ tagged recombinant proteins and purified as described in Wang *et al*. 2015.^15^ AgrC-I and MSP1D1 were expressed and purified as described in Wang *et al*. 2014.^14^

#### Full-length AgrD substrates

AgrD-GyrA-His_7_ fusion proteins were expressed at 8 L scale in BL21(DE3) *E. coli*. For expression, bacteria were grown to an OD_600_ of 0.6 and cooled to 18 °C. Expression was induced by addition of IPTG to 0.5 mM, and bacteria were incubated at 18 °C overnight. Bacteria were harvested through centrifugation, resuspended in a lysis buffer (20 mM Phosphate, 100 mM NaCl, 1 mM PMSF, 1 mM TCEP, pH 7.5) and homogenized by four passes through an Avestin Emulsiflex C3 French press homogenizer (ATA Scientific Instruments) or sonication using an Ultrasonic Liquid Processor Sonic Dismembrator (ThermoFisher). Soluble protein was separated from cellular debris by centrifugation of lysate at 30,000 x g for 30 min at 4 °C. Soluble protein fraction was then loaded onto a pre-equilibrated Ni-NTA resin at a loading of 1 mL resin / L expression volume. Resin was washed with 5 CV wash buffer I (20 mM Phosphate, 500 mM NaCl, 15 mM imidazole, 1 mM TCEP, pH 7.5). All AgrD peptides were then eluted from the Ni-NTA resin by thiol-induced cleavage of the Mxe GyrA intein. For full-length AgrD peptides, an on-column dual thiolysis/hydrolysis was performed. Resin was incubated in 2 CV thiolysis/hydrolysis buffer (20 mM Phosphate, 100 mM NaCl, 50 mM β-mercaptoethanol, 5 mM TCEP, pH 6.9) and 2 CV of elution buffer (20 mM Phosphate, 100 mM NaCl, 6 M GuHCl, pH 6.5). Combined eluant was then purified by RP-HPLC and the peptides characterized by ESI-MS. *AgrD (1-32) thiolactones*: AgrD(1-32)-GyrA-His_6_ fusion proteins from *agr* groups 1 and III were expressed as above for full-length AgrD and immobilized on Ni-NTA resin. This resin was then incubated in 2 CV MESNa thiolysis buffer (20 mM phosphate, 100 mM NaCl, 100 mM MESNa, 5 mM TCEP, pH 6.9), stirring overnight in a round bottom flask under argon at room temperature.

The majority of AgrD-(1-32)-MESNa thioester is insoluble in aqueous buffer, and forms a fine precipitate. AgrD-(1-32)-MESNa thioester, both in solution and as precipitate, was eluted from the column, and the column was washed with 20 CV wash buffer II (20 mM phosphate, 100 mM NaCl, 1 mM TCEP, pH 6.9). The eluent and wash flow-through were combined and centrifuged at 4,000 x g for 10 min at 4 °C to pellet insoluble AgrD-(1-32)-MESNa thioester, which was washed in 2 CV wash buffer II and centrifuged again under the same conditions. The Ni-NTA resin was then treated with 2 CV of elution buffer (20 mM phosphate, 100 mM NaCl, 6 M GuHCl, pH 6.5) and this eluant combined with the pelleted AgrD-(1-32)-MESNa thioester. Solubilization of the pellet in this buffer (which importantly contains no thiols) led to spontaneous cyclization to form the desired AgrD-(1-32)-thiolactone. The cleaved Mxe-GyrA-His_6_ side-product was removed using a reverse Ni-NTA column, and the desired peptide further purified by C18 analytical RP-HPLC. Purified AgrD-(1-32)-thiolactone peptides were lyophilized and stored in DMSO at -20°C.

AgrD-II-(1-32)-thiolactone was made by Boc-solid phase peptide synthesis (SPPS), leveraging the general strategy for the synthesis of *agr* peptides containing C-terminal thiolactones presented in Lyon *et al*. 2002 and Johnson *et al*. 2015.^25, 41^ Peptide synthesis was performed with a 4-methylbenzhydrylamine-copoly(styrene-1% DVB) (MBHA resin) support, pre-functionalized with S-trityl mercaptopropionic acid to form a C-terminal thioester linker. Standard Boc-SPPS was performed, using in situ neutralization and HBTU activation of Boc/Bzl-protected amino acids. After elongation, global deprotection was performed by HF-cleavage and crude linear peptide was precipitated in cold Et_2_O. After lyophilization, linear AgrD (1-32) thioester was cyclized in 200 mM phosphate buffer (pH 7). AgrD-II-(1-32) thiolactone product was then purified by RP-HPLC and characterized by ESI-MS.

### Synthesis of DABCYL-SceD-EDANS and AIPs

#### DABCYL-SceD-EDANS

The fluorescent peptide substrate for SpsB, designated DABCYL-SceD-EDANS^31^ was synthesized by standard Fmoc-SPPS, using an EDANS NovaTag resin (Novabiochem.) Synthesis was performed with 5 eq. of standard N-terminal Fmoc amino acids activated with PyBOP and HOBT (4.9 eq) in DMF. Fmoc deprotection was performed with 4% DBU in DMF. N-terminal DABCYL addition was performed overnight using DABCYL-OSu and DIEA at 2.5 eq. Final deprotection and cleavage was performed using a 95% TFA (v/v), 2.5% TIPS (v/v) and 2.5% water (v/v) cleavage cocktail. TFA was removed by N_2(g)_ and peptide products were precipitated with cold diethyl ether. Precipitated crude peptide was dissolved in 80% HPLC Solvent A/20% HPLC Solvent B and purified by preparative scale RP-HPLC, lyophilized, and taken up in DMSO. Peptide products were characterized by HPLC and LC/MS.

#### AIPs

AIP-I, AIP-II, AIP-III were synthesized as previously reported in Wang *et al*. 2015.^15^ AIP-I-II and AIP-II-I were generated by standard Fmoc-SPPS, on a hydrazine derivatized trityl-ChemMatrix resin (Biotage). Linear hydrazide peptides were oxidized with NaNO_2_, after which thiolysis was performed with MESNa. AIP-MESNa thioesters were purified by RP-HPLC and cyclized in PBS with TCEP at pH 7. Cyclized products were purified again by RP-HPLC, lyophilized, and taken up in DMSO. AIP-I-II and AIP-II-I products were characterized by HPLC and LC/MS.

### *S. aureus* Growth conditions

To initiate each growth experiment, *S. aureus* strains were grown from glycerol stocks at 37 °C overnight in CYGP medium without glucose (10 g/L Casamino acids, 10 g/L yeast extract, 5.9 g/L NaCl, and 60 mM β-glycerophosphate). For strains transduced with *pCN51*, media was supplemented with 10 μg/ml erythromycin and 2 μM CdCl_2_ to select for positive transductants and induce plasmid expression respectively. All strains were prepared for growth curves by subculturing overnight bacterial growths from glycerol stocks, diluting bacteria 1:50 with fresh CYGP medium without glucose and growing diluted cultures for 2 hours. Subcultures were then centrifuged at 8,000 x g, washed twice in sterile PBS, and diluted to an OD_600_ = 0.05 in fresh CYGP medium without glucose. Strains were grown for 24-48 hours at 37 °C in a shaker incubator at 75 mL in sterile 250 mL Erlenmeyer flasks. For growth assays, samples were taken every 1-4 hours for growth monitoring by OD_600_ and AIP production measurement. AIP production was quantified by LC/MS and validated through β-lactamase reporter cell assay when applicable.

### Proteoliposome assembly

The protocol for proteoliposome assembly was adapted from Wang *et al*. 2015.^15^ Proteoliposomes were formed by coincubation of integral membrane proteins with polar lipids in detergent containing buffer, followed by removal of detergent through absorption by BioBeads SM-2 (BioRad) to force the formation of proteoliposomes. Typical setups were performed at 500 μL scale, incubating 5 μM of SEC purified integral membrane protease (MroQ, SpsB, AgrB, etc.) solubilized in detergents with a lipid mixture of 16.25 mM 3:1 mixture of POPC (1-palmitoyl-2-oleoyl-sn-glycero-3-phosphocholine) and POPG (1-palmitoyl-2-oleoyl-sn-glycero-3-phospho-(1’-rac-glycerol)) solubilized in stock vials at 50 mM in 20 mM HEPES buffer containing 150 mM sodium cholate detergent. Phosphate buffer (pH 7.5) or Tris-HCl buffer (pH 7.5) was added to the solution to a final concentration of 30 mM, and water was added to a final volume of 500 μL per setup. Reagents were mixed and allowed to equilibrate at room temperature for 30 minutes on a nutator. After equilibration, activated Bio-Beads SM-2 resin was added to the mixture at 50x w/w relative to the mass of detergent in the assembly solution. The mixture was incubated on Bio-Beads for 2 hours at RT or overnight at 4°C on a nutator. The resulting proteoliposomes were eluted from Bio-Beads and isolated by centrifugation. Final concentration of protein in proteoliposomes was measured by A_280_ relative to empty liposomes assembled by the same protocol.

### Biochemical assays using proteoliposomes

SceD and AgrD-(1-32) thiolactone substrates, solubilized in DMSO, were added to a concentration of 0.5 μM into a solution containing 2 μM of membrane protease in proteoliposomes in 30 mM phosphate, 2.5 mM TCEP, pH 7.5. M131 (SpsB inhibitor) was added to a concentration of 2 μM to relevant reaction mixtures. Full-length AgrD substrates, also solubilized in DMSO, were added to a concentration of 10 μM into a solution containing 5 μM assembled MroQ proteoliposomes and 10 μM assembled cogenetic AgrB proteoliposomes. Empty liposomes, assembled by the same method as proteoliposomes, were added to each reaction mixture to dilute proteoliposomes to target concentrations. To each reaction type, TCEP was added to 2.5 mM, and DMSO added to a final concentration of 1%, with a typical reaction volume of 120 μL. FRET-based cleavage assays were incubated and analyzed in a Spectramax M3 microplate reader (Molecular Devices) at 37 °C for 3 hours, monitoring fluorescence at an excitation of 340 nm and an emission of 510 nm. Proteoliposome-based cleavage assays were incubated at 37 °C for 3-24 hours in a shaker-incubator. Control reactions, performed with empty liposomes and a serial dilution of synthetic AIP-I, -II, or -III, were performed in parallel to generate standard curves for LC-MS analysis. Reactions were stopped by snap freezing or by acidification with neat TFA for solid phase extraction (SPE) purification.

### Solid Phase Extraction Purification of AIPs or Cleavage Products

Sep-PaK Vac 1cc (50 mg) C18 SPE columns (Waters) were activated with 3x 1 mL HPLC solvent B (90% acetonitrile, 10% H_2_O, 0.1% TFA) and equilibrated with 1 mL HPLC solvent A (H_2_O, 0.1% TFA). Analyte-containing solutions, derived from biochemical assays or bacterial growth media, were acidified by addition of TFA to 0.1% and normalized with the addition of an internal standard (fluorescein labeled AIPs, FAM-AIP-I or FAM-AIP-II) to 0.05 μM prior to addition to the SPE column. Analyte solution was allowed to drain and washed with 1 mL 90% HPLC solvent A / 10% HPLC solvent B. AIPs and other cleavage products were eluted from SPE columns with 1 mL 30% HPLC solvent A / 70% HPLC solvent B. UPLC-grade solvent used to generate Solvent A and Solvent B used for wash and elution steps. Eluent containing the compounds of interest was lyophilized overnight.

### LC/MS analysis of protease activity or AIP production

Lyophilized SPE eluent containing AIPs or cleavage products of interest was resuspended in 50% MeCN/0.1% FA in water prior to dilution and analysis by LC/MS. LC/MS analysis was performed on a 6546 High-Resolution Q-TOF (Agilent), utilizing an Atlantis dC_18_ C18 UPLC column (Waters), using a 15-85% gradient in a MeCN/H_2_O solvent system. Extracted ion chromatograms, selecting the mass of each compound of interest and the internal standard, were generated for each analyzed sample. Peptide identity was confirmed by MS-MS and quantified via standard curves generated using purified synthetic AIP standards.

### β-lactamase reporter cell assay

*agr* activation by endogenously produced AIP was detected using a β-lactamase reporter cell assay as previously reported.^25, 42^ Reporter cell lines in *agr* group-I, *agr* group-II, and *agr* group-III backgrounds (see Table 1) were grown at 37°C overnight in CYGP medium without glucose supplemented with 10 μg/ml erythromycin. 100 μL overnight culture was re-inoculated in 10 mL of fresh CYGP media and grown to OD_600_ = 0.6 at 37°C. Media samples taken from growth curve experiments of analyzed *S. aureus* strains were diluted 1:10 in fresh CYGP media and mixed 1:1 with reporter cell lines to a total volume of 100 μL. Reporter cells were then grown at 37°C for one hour in a Multiskan GO 96-well plate reader and shaker incubator (ThermoFisher), monitoring growth of reporter cells over time at OD_650_. Cells were then diluted 1:1 with 200 μM nitrocefin prepared in 100 mM phosphate, 5 mM NaN_3,_ and 20% propylene glycol, pH 6.0, to a total volume of 100 μL in a fresh 96 well plate. Gain of absorbance signal from β-lactamase activity on nitrocefin was measured over time at 490 nm, incubating the reaction mixture at 37°C for 45 minutes with shaking in the plate reader.

### Nanodisc assembly

Reconstitution of purified recombinant AgrC-I into lipid nanodiscs was performed as previously described in Wang *et al*. 2014, utilizing MSP1D1 membrane scaffold protein.^14^

### One-pot AIP biosynthesis and AgrC autokinase assay

Reactions were performed by combining proteoliposomes (indicated combinations of SpsB, AgrB-I, AgrB-II and MroQ) with AgrC-I nanodiscs in a buffer containing 50 mM Tris-HCl, 15 mM HEPES, 100 mM NaCl, 10 mM MgCl_2_, 1 mM TCEP. The final concentration of each protease was 1.7 μM, whereas AgrC-I was 0.7 μM. To this mixture was added AgrD-I to a final concentration of 40 μM. Some reactions also contained AIP-I (10 μM) and/or the AgrC inhibitor^36^ QQ-3 (5 μM). Note, addition of the AIP-I positive control allowed us to establish a theoretical maximum of *agr* activation under equivalent assay conditions. Total DMSO concentration was brought to 2.5% in all assays, consistent with precedent.^14, 16, 17, 36^ Prior to addition of ATP-γ-^32^P, aliquots of each reaction setup were removed for loading control analysis by SDS-PAGE/CBB stain. ATP was then added to the reaction mixture to final concentrations of 10 μM cold ATP and 10 μM ATP-γ-^32^P (10 Ci/mmol) (PerkinElmer). Reactions were incubated for 40 minutes at 37 °C. The reactions were quenched by the addition of SDS-PAGE loading dye and reaction components were resolved by SDS-PAGE using a 12% Tris-HCl gel. Gels processed by drying and exposure to Carestream Kodak Biomax CAR film (Sigma Aldrich) for detection of ^32^P phosphorylation of AgrC-I. Imaging was performed on an Amersham ImageQuant 800 (Cytiva). Densitometry analysis was performed by ImageJ.

