## Supplemental Figures for "Reconstitution of the *S. aureus agr* quorum sensing pathway reveals a direct role for the integral membrane protease MroQ in pheromone biosynthesis"

**Figure S1. In vitro activity of SpsB. A**. SDS-PAGE analysis of purified recombinant $His_5$-TEV-SpsB. **B**. SDS-PAGE analysis of recombinant $His_5$-TEV-SpsB reconstituted into POPC:POPG 1:3 proteoliposomes. **C**. RP-HPLC chromatogram and ESI-MS spectrum of DABCYL-SceD-EDANS. **D.** Initial rate of gain of fluorescence from cleavage of fluorogenic SpsB substrate DABCYL-SceD-EDANS by empty liposomes (EL) or SpsB 1:3 proteoliposomes. Data presented as the mean ± s.d. (n=3 biological replicates). **E.** EIC traces of DABCYL-SceD-EDANS SpsB cleavage reaction in the presence or absence of M131 inhibitor. The cleavage product SET-EDANS is only observed in the absence of inhibitor.

**Figure S2. Characterization of AgrD-(1-32)-thiolactone substrates**. **A**. RP-HPLC and ESI-MS of AgrD-I (1-32) thiolactone. **B**. RP-HPLC and ESI-MS of AgrD-II (1-32) thiolactone. **C**. RP-HPLC and ESI-MS of AgrD-III (1-32) thiolactone.

**Figure S3. Characterization of synthetic AIP standards. A**. RP-HPLC and ESI-MS of AIP-I. **B**. RP-HPLC and ESI-MS of AIP-II. **C**. RP-HPLC and ESI-MS of AIP-III.

**Figure S4. In vitro SpsB activity assays**. **A**. SpsB proteoliposomes were mixed with indicated AgrD (1-32) thiolactones in the presence of the fluorogenic DABCYL-SceD-EDANS substrate and optionally M131 inhibitor. Cleavage of the fluorescence substrate was monitored by fluorescence (510 nm) over time. Data presented as the mean ± s.d. (n=3 biological replicates). **B**. Growth of WT, Δ*cro/cI*, and Δ*cro/cI*/Δ*spsB* variants of group-I *S. aureus* over 24 hours, monitored by $OD_{600}$ . Data presented as the mean ± s.d. (n=3 biological replicates). **C**. Standard curve for the quantification of AIP-I by LC-MS. Data presented as the mean ± s.d. (n=3 replicates). **D**. Quantified AIP-I production by WT, Δ*cro/cI*, and Δ*cro/cI*/Δ*spsB* variants of group-I *S. aureus* over 24 hours, determined by LC-MS. Data presented as the mean ± s.d. (n=3-4 biological replicates).

**Figure S5. Effect of MroQ knockout on AIP production in *S. aureus* groups 1-III. A**. Standard curve for the quantification of AIP-I, AIP-II, and AIP-III relative to internal standard by LC-MS. Data presented as the mean ± s.d. (n=3 replicates). **B**. Quantified AIP production by WT + empty vector, Δ*mroQ* + empty vector, Δ*mroQ* + *mroQ*, and Δ*mroQ* + *mroQ*$^{mut}$ variants of group-I, -II, and -III *S. aureus* over 48 hours, determined by LC-MS. Data presented as the mean ± s.d. (n=3 replicates). **C**. Relative AIP production of WT and Δ*mroQ* variants of group-I, -II, and -III *S. aureus* as determined by reporter cell assay. Data presented as the mean ± s.d. (n=3 replicates).

**Figure S6. MroQ catalytic activity is responsible for AgrD (1-32) thiolactone cleavage in group-I and group-I AIP maturation**. **A**. SDS-PAGE analysis of recombinant MBP-MroQ-$His_6$ and $His_6$-MBP-$MroQ^{mut}$-HA. Identity of proteins was confirmed by tryptic digestion and LC-MS (data not shown). **B**. MroQ or $MroQ^{mut}$ proteoliposomes were mixed with indicated AgrD (1-32) thiolactone intermediates. Reaction mixtures containing a spike-in internal standard were then subjected to a SPE step and analyzed by LC-MS. Shown are the EIC traces for the expected AIP products and the internal standard. A synthetic AIP treated with empty liposomes served as a positive control (black trace). **C**. AIP-III levels produced by MroQ, $MroQ^{mut}$ or SpsB proteoliposomes were quantified by LC-MS using a standard curve approach employing synthetic AIP-III. Data presented as the mean ± s.d. (n=3 biological replicates). **D.** LC-MS/MS of AIP-III produced by cleavage of AgrD-III (1-32) thiolactone by MroQ proteoliposomes (blue) and synthetic AIP-III (black).

**Figure S7. Characterization of AgrB and AgrC.** **A**. SDS-PAGE analysis of recombinant AgrB-I-$His_6$ and AgrB-II-$His_6$ reconstituted into proteoliposomes. Note, AgrB-I runs primarily as a dimer, which is consistent with a previous report.[15] **B**. SDS-PAGE analysis of recombinant AgrC-I reconstituted into lipid nanodiscs. MSP, membrane scaffold protein.

**Figure S8. Multiple sequence alignment of AgrD-I, AgrD-II, and AgrD-III.** Peptide domain architecture indicated above. Cleavage sites indicated in red dashed lines for the formation of AIP-I, AIP-II, and AIP-III respectively.

**Figure S9. Characterization of chimeric AgrD (1-32) thiolactones and chimeric AIP products**. **A**. RP-HPLC and ESI-MS analysis of AgrD-I-II-I (1-32) thiolactone and AgrD-II-I-II (1-32) thiolactone. **B**. RP-HPLC and ESI-MS analysis of AIP-I-II and AIP-II-I.

**Figure S10. Overview of *S. aureus* chimeric AgrD mutant AIP production assays**. Constitutively active AgrC activates AgrA for upregulation of expression of AgrD chimeras. The

C-terminus of AgrD chimeras are processed normally to yield chimeric biosynthetic intermediates. These intermediates are then cleaved and secreted into the extracellular milieu. Putative cleavage products shown based on position of N-terminal cleavage.

#### Supplementary Table 1

| Strain | Description | Reference |
|---|---|---|
| RN1 | Group-I$^{wild\ type}$ strain | Peng *et al.* 1988 |
| RN1$^{\Delta cro/cI}$ | Group-I$^{\Delta cro/cI}$ strain, knockout of *cro/cI.* | This study |
| RN1$^{\Delta cro/cI/\Delta spsB}$ | Group-I$^{\Delta cro/cI/\Delta spsB}$ strain, knockout of s*psB* and *cro/cI*. | This study |
| RN1$^{\Delta mroQ}$ | Group-I$^{\Delta mroQ}$ strain, knockout of *mroQ* | This study |
| RN6607$^{wild\ type}$ | Group-II$^{wild\ type}$ strain. | Ji et al. 1997 |
| RN6607$^{\Delta mroQ}$ | Group-II$^{\Delta mroQ}$ strain, knockout of *mroQ* | This study |
| MW2$^{wild\ type}$ | Group-III$^{wild\ type}$ strain. | CDC, 1999 |
| MW2$^{\Delta mroQ}$ | Group-III$^{\Delta mroQ}$ strain, knockout of *mroQ* | This study |
| RN9222 | Reporter strain for the identification of AIP-I in growth media by β-lactamase expression and nitrocefin assay | Lyon *et al.* 2000 |
| RN9367 | Reporter strain for the identification of AIP-II in growth media by β-lactamase expression and nitrocefin assay | Lyon *et al.* 2000 |
| RN9532 | Reporter strain for the identification of AIP-III in growth media by β-lactamase expression and nitrocefin assay | Lyon *et al.* 2002 |
| RN7206 | *agr*-null strain | Novick *et al.* 1993 |
| RN4220 | Restriction-deficient *S. aureus* host strain | Kreiswirth *et al.* 1983 |
| TMD1211 | AgrD-I-II-I chimera strain. *agr*-null (RN7206) strain with chromosomal integration of *agrB-I-agrD-I-II-I-agrC*$^{R238K}$*-agrA* at *attC* site. | This study |
| TMD2122 | AgrD-II-I-II chimera strain. *agr*-null (RN7206) strain with chromosomal integration of *agrB-II-agrD-II-I-II-agrC*$^{R238K}$*-agrA* at *attC* site. | This study |

**Supplementary Table 2**

| Plasmid Name | Description | Reference |
|---|---|---|
| pD1211 | pJC1111 encoding SaPI1 attC:: *agrP2-agrB-I-agrD-I-II-I-agrC*$^{R238K}$*-agrA* for chromosomal integration of AgrD-I-II-I chimera. | This study |
| pD2122 | pJC1111 encoding SaPI1 attC::*agrP2-agrB-II-agrD-II-I-II-agrC*$^{R238K}$*-agrA* for chromosomal integration of AgrD-II-I-II chimera. | This study |
| pMroQ | pET24b encoding *MBP-mroQ-His*$_6$ for the expression MBP-MroQ | This study |
| pMroQ$^{mut}$ | pET24b encoding *His*$_6$*-MBP-mroQ-HA* for the expression MBP-MroQ$^{mut}$ | This study |
| pBW-B1-2 | pET24b encoding *agrB-I-His*$_6$ for the expression of AgrB-I | Wang *et al.* 2015 |
| pBW-B2-2 | pET24b encoding *agrB-II-His*$_6$ for the expression of AgrB-II | Wang *et al.* 2015 |
| pBW-C1-1 | pET24b encoding *agrC-I-His*$_6$ for the expression of AgrC-I | Wang *et al.* 2014 |
| pMSP1D1 | pET15b encoding *His*$_6$*-Thrombin-MSP1D1* for the expression of MSP1D1. | This study |
| pJJ60 | pET15b encoding *His*$_5$*-TEV-spsB* for the expression of SpsB | This study |
| pBW-D1-8 | Modified pTXBI encoding *agrD-I-(1-32)-gyrA-His*$_7$ for the production of AgrD-I (1-32) thiolactone | Wang *et al.* 2015 |
| pBW-D3-8 | Modified pTXBI encoding *agrD-III-(1-32)-gyrA-His*$_7$ for the production of AgrD-III (1-32) thiolactone | Wang *et al.* 2015 |
| pBW-D1-1 | Modified pTXBI encoding *agrD-I-gyrA-His*$_7$ for the production of AgrD-I | Wang *et al.* 2015 |
| pBW-D2-1 | Modified pTXBI encoding *agrD-II-gyrA-His*$_7$ for the production of AgrD-II | Wang *et al.* 2015 |
| pDI-II-ITL | Modified pTXBI encoding *agrD-I-II-I-(1-32)-gyrA-His*$_7$ for the production of AgrD-I-II-I (1-32) thiolactone | This study |
| pDII-I-IITL | Modified pTXBI encoding *agrD-II-I-II (1-32)-gyrA-His*$_7$ for the production of AgrD-II-I-II (1-32) thiolactone | This study |
| pCN51 | pCN51 $Cd^{2+}$-inducible plasmid | Charpentier *et al.* 2004 |
| pMroQ$_{comp}$ | pCN51 encoding *mroQ* for genetic complementation | This study |
| pMroQ$^{mut}_{comp}$ | pCN51 encoding *mroQ*$^{mut}$ for genetic complementation control | This study |

### Figure S1

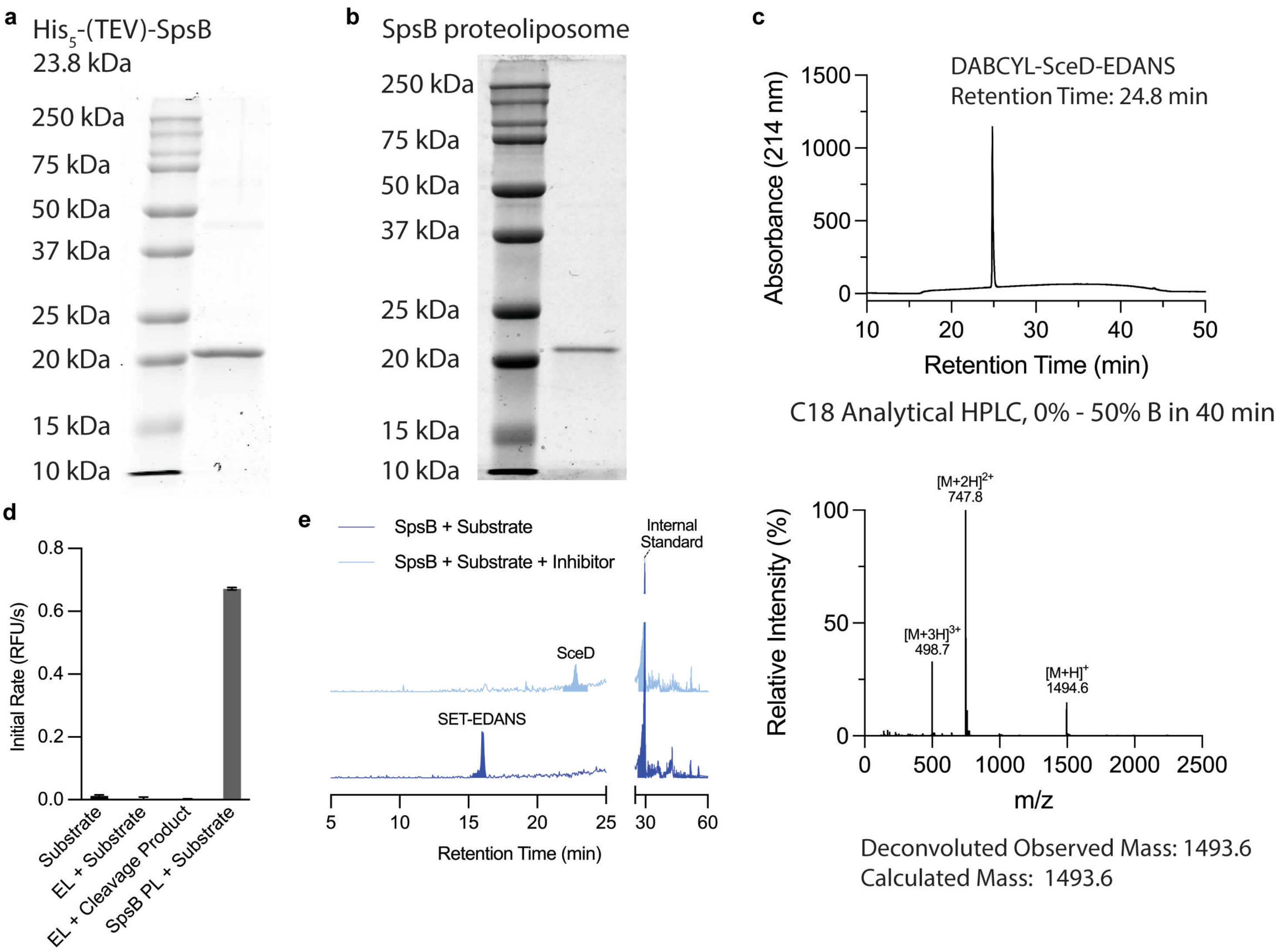

a
His5-(TEV)-SpsB
23.8 kDa
250 kDa
75 kDa
50 kDa
37 kDa
25 kDa
20 kDa
15 kDa
10 kDa
b
SpsB proteoliposome
250 kDa
75 kDa
50 kDa
37 kDa
25 kDa
20 kDa
15 kDa
10 kDa
c
DABCYL-SceD-EDANS
Retention Time: 24.8 min
Absorbance (214 nm)
Retention Time (min)
C18 Analytical HPLC, 0% - 50% B in 40 min
[M+2H]2+
747.8
[M+3H]3+
498.7
[M+H]+
1494.6
Relative Intensity (%)
m/z
Deconvoluted Observed Mass: 1493.6
Calculated Mass: 1493.6
d
Initial Rate (RFU/s)
Substrate
EL + Substrate
EL + Cleavage Product
SpsB PL + Substrate
e
SpsB + Substrate
SpsB + Substrate + Inhibitor
Internal Standard
SceD
SET-EDANS
Retention Time (min)

### Figure S2

**a**

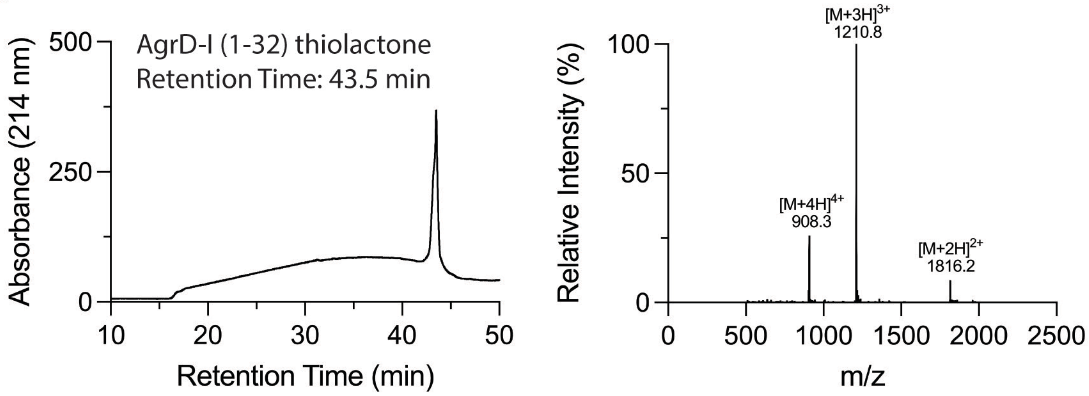


C18 Analytical HPLC, 0% - 100% B in 30 min

Deconvoluted
Observed Mass: 3629.4 Da
Calculated Mass: 3628.8 Da

**b**

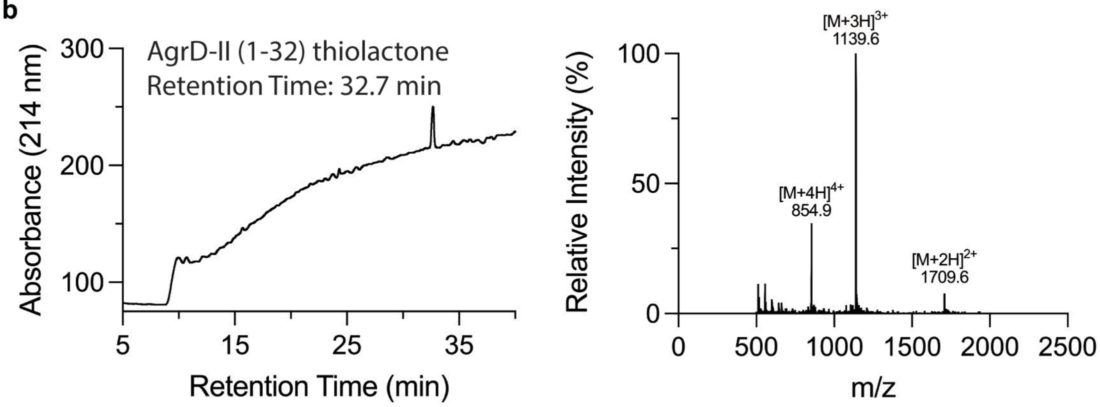


C18 LC-MS, 0% - 90% B in 35 min

Deconvoluted
Observed Mass: 3414.8 Da
Calculated Mass: 3414.8 Da

**c**

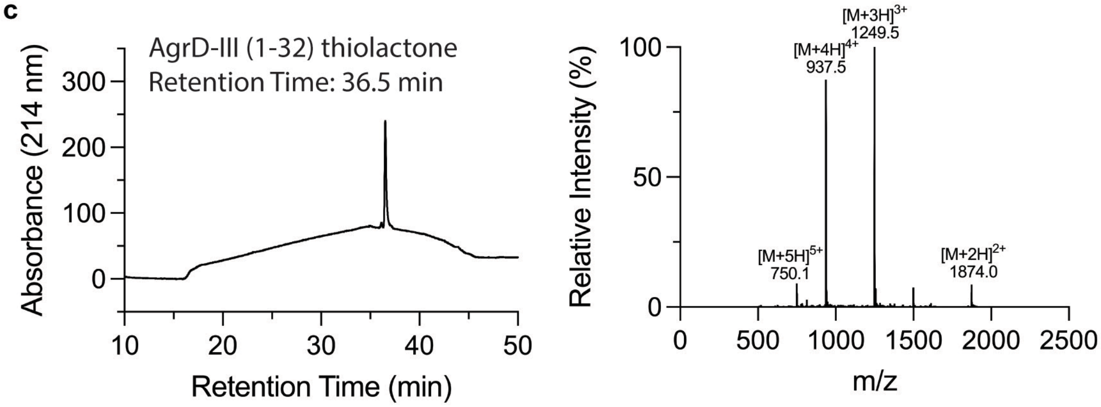


C18 Analytical HPLC, 0% - 100% B in 30 min

Deconvoluted
Observed Mass: 3745.0 Da
Calculated Mass: 3745.0 Da

### Figure S3

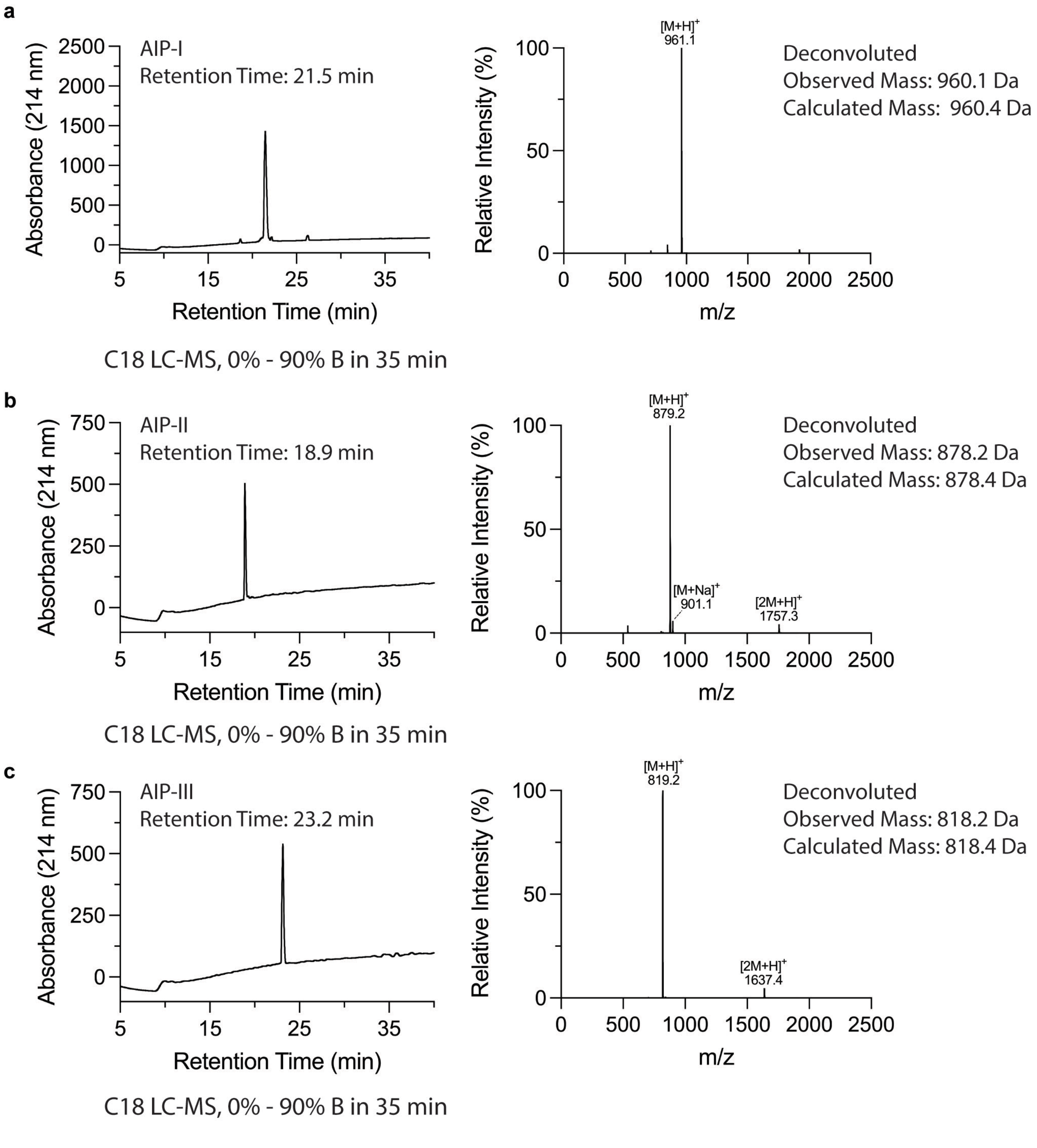
a
AIP-I
Retention Time: 21.5 min
Absorbance (214 nm)
Retention Time (min)
C18 LC-MS, 0% - 90% B in 35 min
[M+H]+
961.1
Relative Intensity (%)
m/z
Deconvoluted
Observed Mass: 960.1 Da
Calculated Mass: 960.4 Da
b
AIP-II
Retention Time: 18.9 min
Absorbance (214 nm)
Retention Time (min)
C18 LC-MS, 0% - 90% B in 35 min
[M+H]+
879.2
[M+Na]+
901.1
[2M+H]+
1757.3
Relative Intensity (%)
m/z
Deconvoluted
Observed Mass: 878.2 Da
Calculated Mass: 878.4 Da
c
AIP-III
Retention Time: 23.2 min
Absorbance (214 nm)
Retention Time (min)
C18 LC-MS, 0% - 90% B in 35 min
[M+H]+
819.2
[2M+H]+
1637.4
Relative Intensity (%)
m/z
Deconvoluted
Observed Mass: 818.2 Da
Calculated Mass: 818.4 Da

### Figure S4

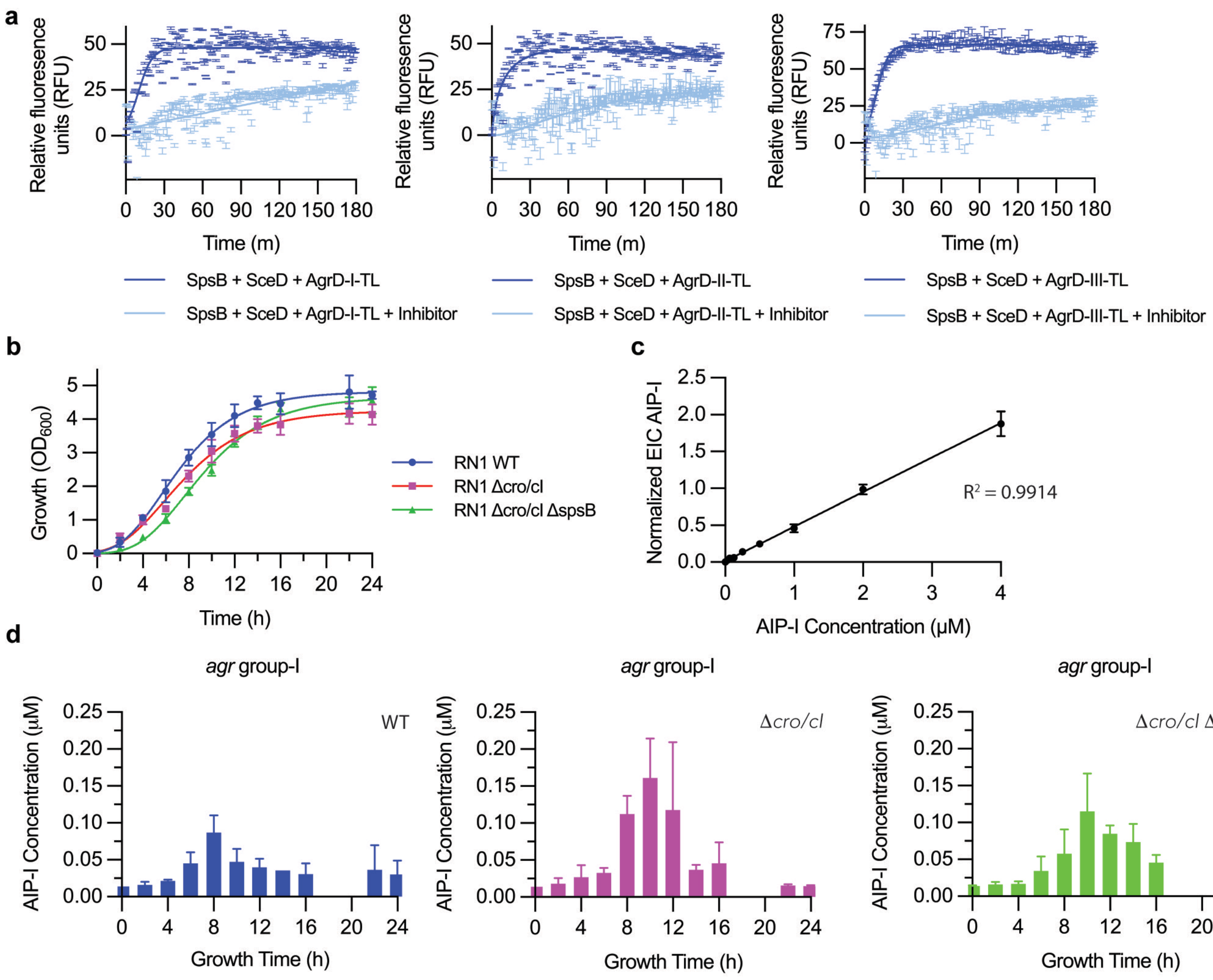

a
Relative fluoresence units (RFU)
Time (m)
SpsB + SceD + AgrD-I-TL
SpsB + SceD + AgrD-I-TL + Inhibitor
SpsB + SceD + AgrD-II-TL
SpsB + SceD + AgrD-II-TL + Inhibitor
SpsB + SceD + AgrD-III-TL
SpsB + SceD + AgrD-III-TL + Inhibitor
b
Growth (OD600)
Time (h)
RN1 WT
RN1 Δcro/cI
RN1 Δcro/cI ΔspsB
c
Normalized EIC AIP-I
AIP-I Concentration (μM)
R2 = 0.9914
d
agr group-I
AIP-I Concentration (μM)
Growth Time (h)
WT
Δcro/cI
Δcro/cI ΔspsB

### Figure S5

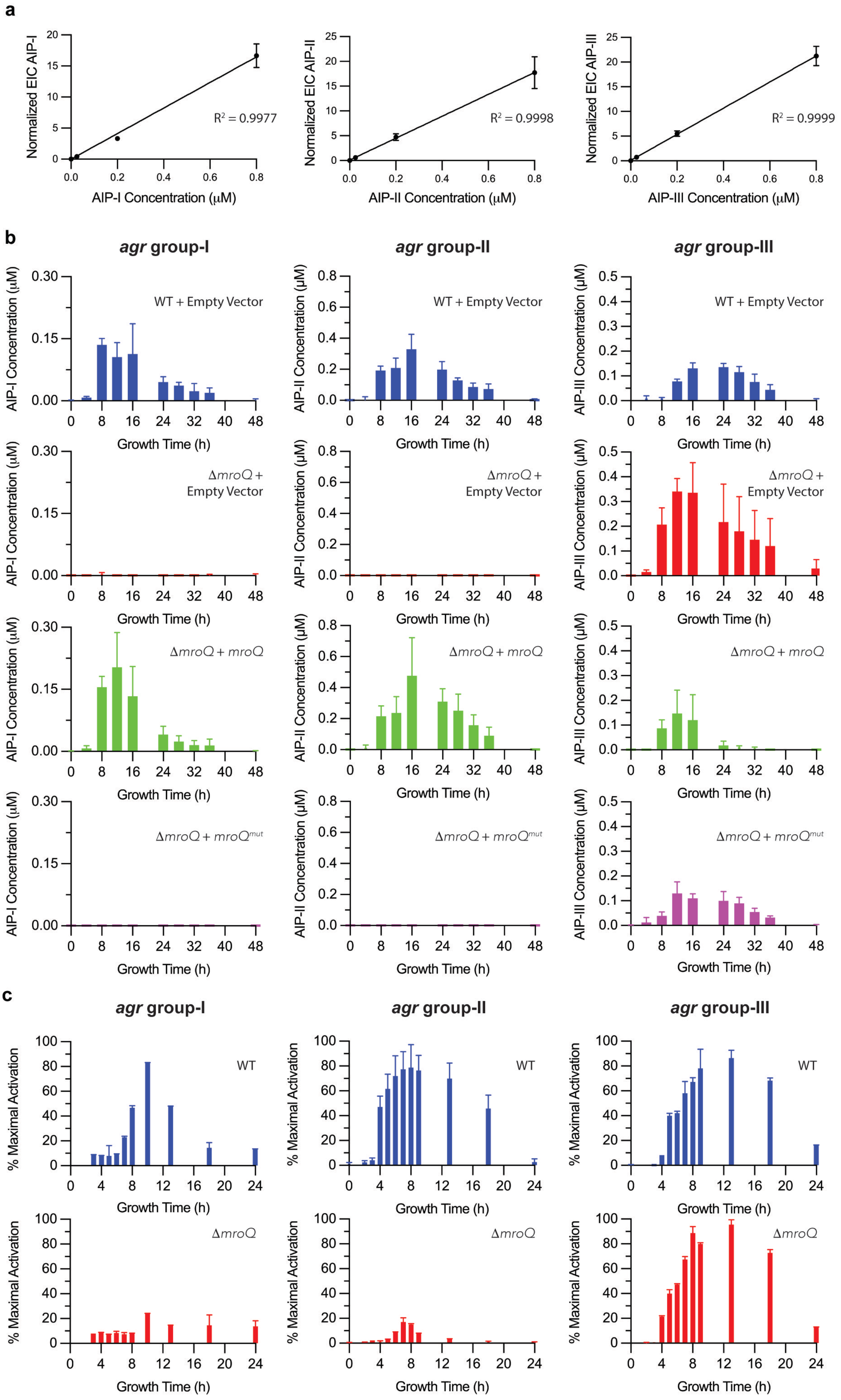

a
Normalized EIC AIP-I
AIP-I Concentration (μM)
R² = 0.9977
Normalized EIC AIP-II
AIP-II Concentration (μM)
R² = 0.9998
Normalized EIC AIP-III
AIP-III Concentration (μM)
R² = 0.9999
b
agr group-I
agr group-II
agr group-III
AIP-I Concentration (μM)
AIP-II Concentration (μM)
AIP-III Concentration (μM)
Growth Time (h)
WT + Empty Vector
ΔmroQ + Empty Vector
ΔmroQ + mroQ
ΔmroQ + mroQmut
c
agr group-I
agr group-II
agr group-III
% Maximal Activation
Growth Time (h)
WT
ΔmroQ

### Figure S6

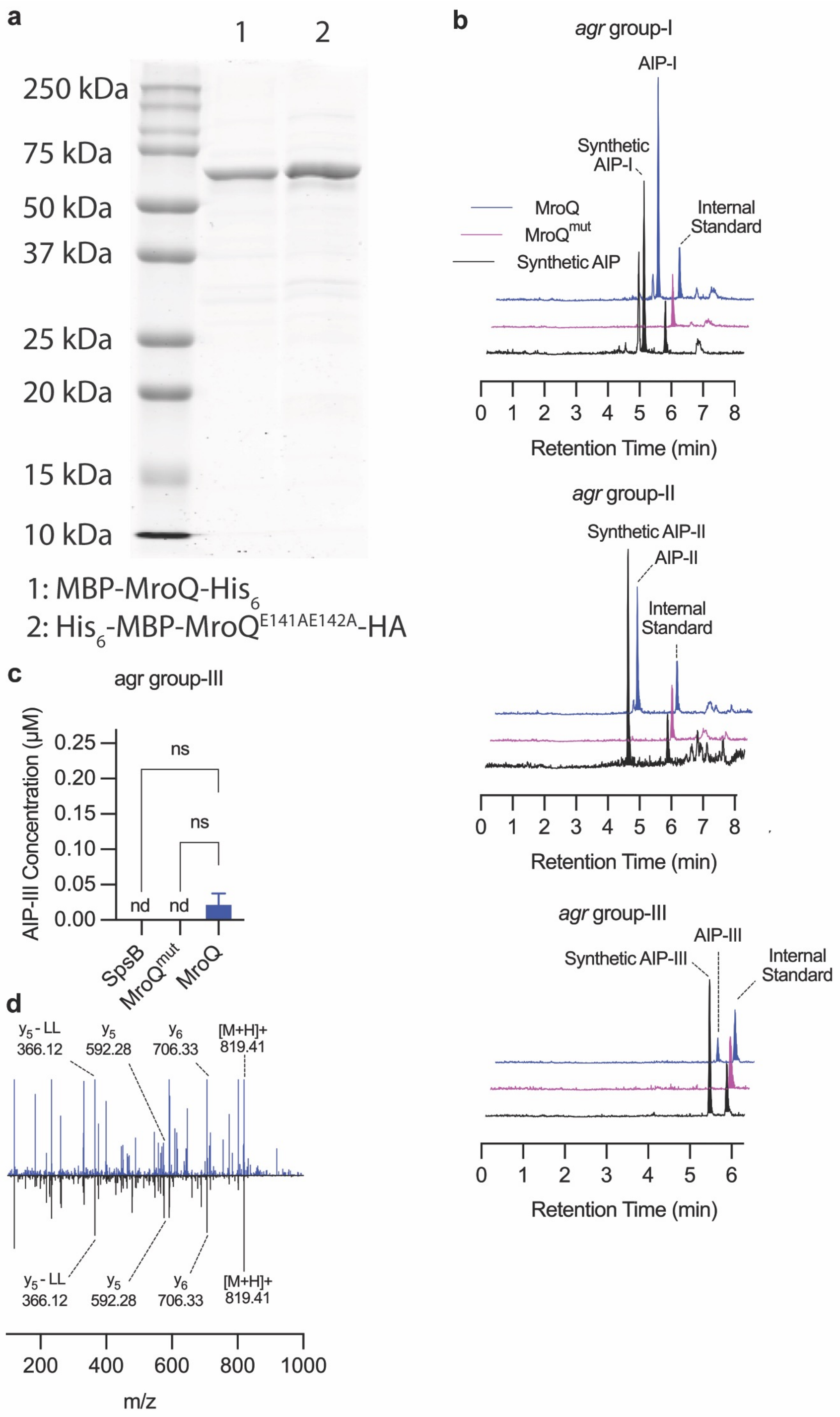

a
1
2
250 kDa
75 kDa
50 kDa
37 kDa
25 kDa
20 kDa
15 kDa
10 kDa
1: MBP-MroQ-His6
2: His6-MBP-MroQE141AE142A-HA
b
agr group-I
AIP-I
Synthetic AIP-I
MroQ
MroQmut
Synthetic AIP
Internal Standard
Retention Time (min)
agr group-II
Synthetic AIP-II
AIP-II
Internal Standard
Retention Time (min)
agr group-III
AIP-III
Synthetic AIP-III
Internal Standard
Retention Time (min)
c
agr group-III
AIP-III Concentration (µM)
ns
ns
nd
nd
SpsB
MroQmut
MroQ
d
y5 - LL 366.12
y5 592.28
y6 706.33
[M+H]+ 819.41
m/z

#### Figure S7

**a**

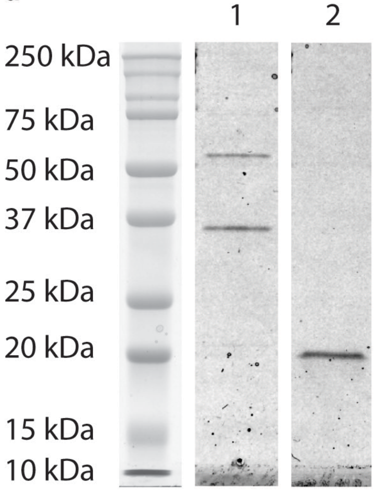


1: AgrB-I-$His_6$ proteoliposome
2: AgrB-II-$His_6$ proteoliposome

**b**

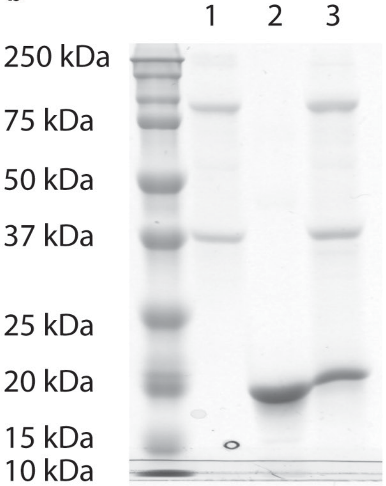


1: AgrC-I-$His_6$
2: MSP1D1
3. AgrC-I-$His_6$/MSP1D1 nanodisc

Figure S8

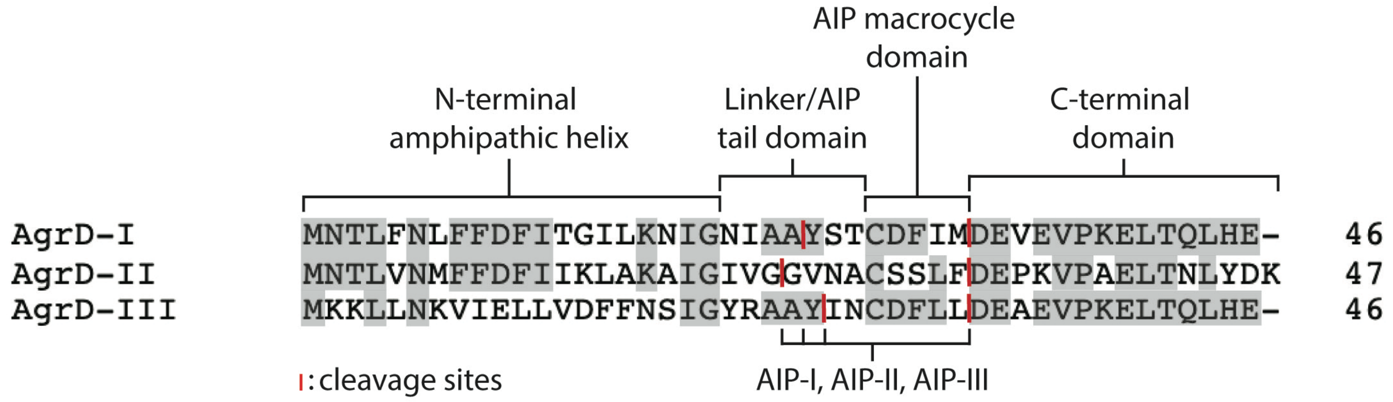
AIP macrocycle domain
N-terminal amphipathic helix
Linker/AIP tail domain
C-terminal domain
AgrD-I MNTLFNLFFDFITGILKNIGNIAAYSTCDFIMDEVEVPKELTQLHE- 46
AgrD-II MNTLVNMFFDFIIKLAKAIGIVGGVNACSSLFDEPKVPAELTNLYDK 47
AgrD-III MKKLLNKVIELLVDFFNSIGYRAAYINCDFLLDEAEVPKELTQLHE- 46
: cleavage sites
AIP-I, AIP-II, AIP-III

Figure S9

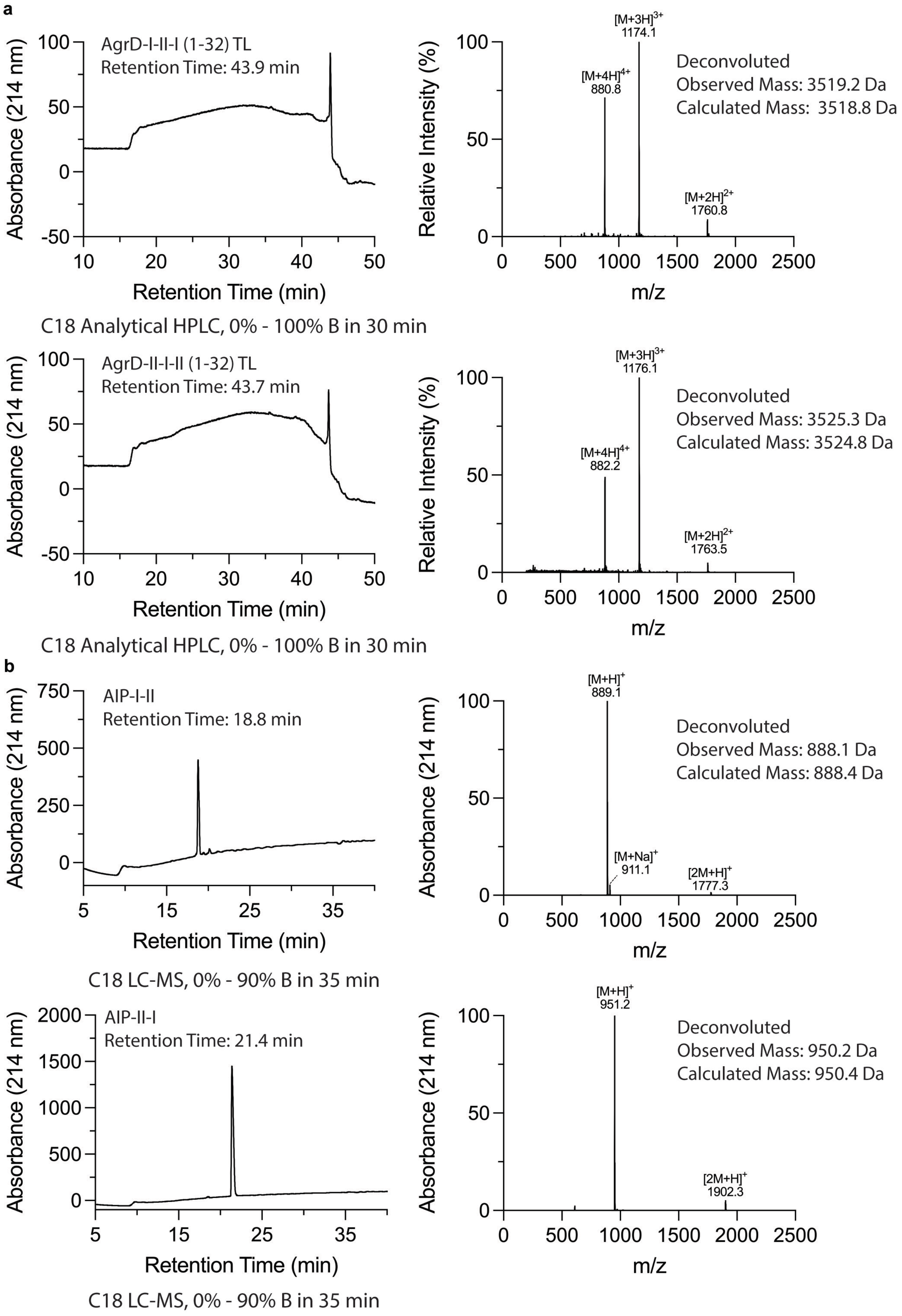
a
AgrD-I-II-I (1-32) TL
Retention Time: 43.9 min
Absorbance (214 nm)
Retention Time (min)
[M+3H]3+ 1174.1
[M+4H]4+ 880.8
[M+2H]2+ 1760.8
Deconvoluted
Observed Mass: 3519.2 Da
Calculated Mass: 3518.8 Da
Relative Intensity (%)
m/z
C18 Analytical HPLC, 0% - 100% B in 30 min
AgrD-II-I-II (1-32) TL
Retention Time: 43.7 min
[M+3H]3+ 1176.1
[M+4H]4+ 882.2
[M+2H]2+ 1763.5
Deconvoluted
Observed Mass: 3525.3 Da
Calculated Mass: 3524.8 Da
C18 Analytical HPLC, 0% - 100% B in 30 min
b
AIP-I-II
Retention Time: 18.8 min
[M+H]+ 889.1
[M+Na]+ 911.1
[2M+H]+ 1777.3
Deconvoluted
Observed Mass: 888.1 Da
Calculated Mass: 888.4 Da
C18 LC-MS, 0% - 90% B in 35 min
AIP-II-I
Retention Time: 21.4 min
[M+H]+ 951.2
[2M+H]+ 1902.3
Deconvoluted
Observed Mass: 950.2 Da
Calculated Mass: 950.4 Da
C18 LC-MS, 0% - 90% B in 35 min

### Figure S10

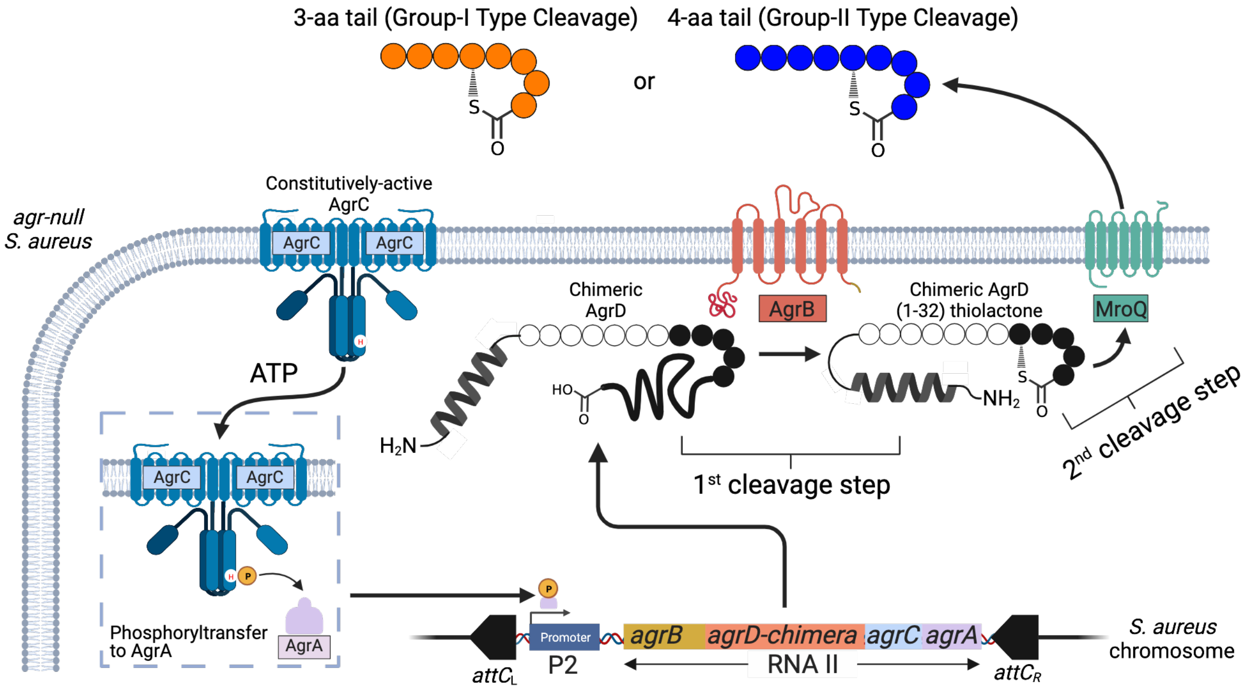

3-aa tail (Group-I Type Cleavage)
or
4-aa tail (Group-II Type Cleavage)
S
O
agr-null
S. aureus
Constitutively-active
AgrC
AgrC
AgrC
ATP
Chimeric
AgrD
AgrB
Chimeric AgrD
(1-32) thiolactone
MroQ
HO
H2N
NH2
1st cleavage step
2nd cleavage step
AgrC
AgrC
Phosphoryltransfer
to AgrA
AgrA
Promoter
P2
agrB
agrD-chimera
agrC
agrA
RNA II
attCL
attCR
S. aureus
chromosome
